# Mapping NMD-coupled alternative splicing in iPSC-derived brain cells: a resource for therapeutic discovery

**DOI:** 10.64898/2026.01.19.700316

**Authors:** Kim N Wijnant, Imke ME Schuurmans, Annika Mordelt, Ka Man Wu, Michael Kwint, Lisenka ELM Vissers, Nael Nadif Kasri

## Abstract

Therapeutic options for neurodevelopmental disorders (NDD) are expanding. Targeting alternative splicing (AS) events linked to nonsense-mediated decay (NMD) offers a promising way to boost gene expression in haploinsufficiency disorders. However, naturally occurring NMD-coupled AS (NMD AS) events in brain cells remain poorly characterized. Here, we integrate long- and short-read RNA sequencing of NMD-inhibited induced pluripotent stem cell-derived excitatory neurons, astrocytes, and microglia to map and prioritize NMD AS events most suitable for therapeutic intervention. We developed an optimized prediction framework and provide an open access, queryable, database cataloging the existence and abundance of NMD AS events across these cell types. Querying this resource, we identified 936 NMD-sensitive AS events in 250 autosomal-dominant NDD genes and nominate 60 NMD AS events in haploinsufficient genes underlying 42 NDDs that are abundant in at least one cell type and thus represent promising therapeutic targets. These included a previously targeted NMD AS event in *SCN1A*, and known NMD AS events in *CHD2, EZH2, and NR4A2*, for which we confirm high cell type-specific abundance, as well as an newly identified abundant event in *PHIP*. Beyond NDD genes, we identify 1,817 differentially spliced AS events including NMD AS events, highlighting the potential functional role of (NMD) AS within brain cell-specific regulatory programs. This framework and resource enables systematic discovery and prioritization of therapeutically targetable NMD AS events and establish a cell-type resolved atlas to guide splice-modulating strategies in NDDs.

## Introduction

Alternative splicing (AS) is a fundamental post-transcriptional mechanism that expands the coding potential of the human genome. AS is especially prominent in the brain, where it contributes to neurodevelopment, neuronal differentiation, and synaptic function (1). While AS generates proteomic diversity, it can also introduce premature stop codons, rendering transcripts sensitive to nonsense-mediated decay (NMD). Like non-NMD-sensitive AS (non-NMD AS), NMD-sensitive AS (NMD AS) may participate in neurodevelopment and synaptic function (2–7).

Beyond its potential physiological roles, NMD AS presents a therapeutic opportunity to upregulate gene expression (8). Antisense oligonucleotides (AONs) can be designed to inhibit NMD AS, a strategy known as targeted augmentation of nuclear gene output (TANGO) (8). This strategy is particularly relevant in disorders caused by dominant pathogenic loss-of-function variants. For these disorders inhibiting NMD AS on the wild-type allele can increase gene expression and compensate for the disease-causing loss-of-function allele. Because AONs, can be delivered directly to the central nervous system via intrathecal administration, TANGO represent a promising modality for neurological disorders, especially monogenic neurodevelopmental disorders (NDDs) (9). In Dravet syndrome (OMIM: 607208), inhibition of an alternative splicing event that triggers NMD in *SCN1A* has been shown to increase *SCN1A* expression, providing proof-of-concept for this therapeutic strategy (10, 11). Additional NDD-associated genes were found to exhibit NMD AS, including *DLG4* (SHINE syndrome, OMIM: 618793), *SYNGAP1* (*SYNGAP1* syndrome, OMIM: 612621), *CHD2* (*CHD2* syndrome, OMIM: 615369), and *PCCA* (Propionicacidemia, OMIM: 606054) (*7, 8, 12, 13*). Disorders linked to these genes may benefit from TANGO-mediated upregulation. Consistent with this idea, inhibition of NMD AS in *SYNGAP1* and *PCCA* has been shown to increase gene output in patient-derived cells (7, 12, 13).

To date, most NMD AS events have been inferred from reference genome annotations (8, 14, 15), an approach that overlooks the temporal- and cell-type specific expression patterns that are critical for therapeutic targeting. A cell-type resolved, experimentally obtained atlas of NMD AS in human brain-relevant cells is therefore essential but is currently lacking. Induced pluripotent stem cell (iPSC)-derived models offer a practical solution by enabling access to otherwise inaccessible, brain-relevant cell types such as neurons, astrocytes, and microglia. In addition, it permits experimental NMD inhibition to estimate abundance of NMD-sensitive transcripts.

Here we systematically assessed the presence and abundance of NMD and non-NMD AS events in iPSCs, as well as in iPSC-derived excitatory neurons, astrocytes, and microglia. We developed and validated two complementary methods for NMD AS identification and quantification by integrating short-read and long-read RNA sequencing in NMD-inhibited cells. Leveraging these methods, we created an open access online, searchable database that catalogs NMD and non-NMD AS events and their abundances in these cell types, providing ample opportunities for further research. Here, we focused on two diverse use cases. First, to deepen our understanding of brain development, we characterize cell-type-specific differential splicing, including NMD-associated programs, thereby delineating therapeutic opportunities and potential liabilities for splice-modulating therapies within brain cell-specific regulatory networks. Second, to inform therapeutic strategies for NDDs, we validate known NMD-associated AS events, discover previously unrecognized events, and quantitatively prioritize candidate targets for TANGO-mediated upregulation.

## Results

### 1. Generation and validation of an NMD AS database

#### 1.1 Identification of NMD-sensitive isoforms and AS events

To establish and validate a comprehensive database of NMD AS events, we first identified NMD-sensitive isoforms and their associated AS events across undifferentiated iPSCs and iPSC-derived mature neurons, microglia, and astrocytes under control (dimethyl sulfoxide, DMSO) and NMD-inhibited (cycloheximide, CHX) conditions (Figure 1A) (16–19). Using HiFi-based Iso-Seq long-read RNAseq we generated 3.6M-11.4M full-length sequence reads per sample with a mean read length ranging from 2,113 to 2,977 bases (Supplementary data 1).

**Figure 1.**
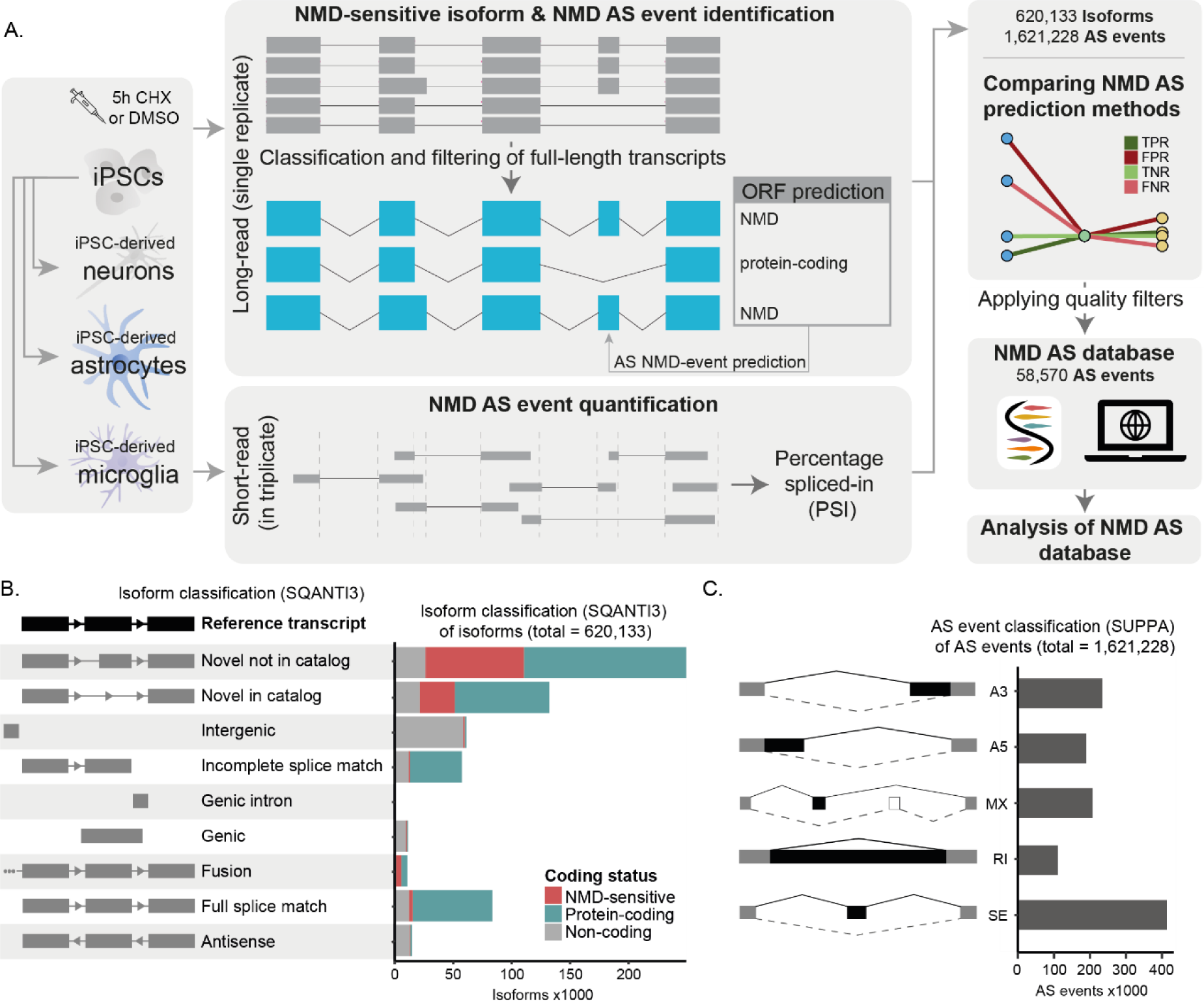
NMD AS database generation and analysis. **A.** Schematic overview of the generation of the NMD AS database using long- and short-read RNAseq data from iPSCs and iPSC-derived cells of the brain. **B.** SQANTI3 isoform classification of all isoforms with graphical explanation of each transcript class compared to an example reference transcript (black). **C.** SUPPA alternative splice event classification of the alternative splice events with graphical explanation of each AS event type. AS = alternative splicing, CHX = cycloheximide, DMSO = dimethyl sulfoxide, iPSC = induced pluripotent stem cell, NMD = nonsense-mediated mRNA decay, ORF = open reading frame, PSI = percentage spliced in, TPR = true positive rate, FPR = false positive rate, TNR = true negative rate, FNR = false negative rate, NMD = nonsense-mediated mRNA decay, A3 = alternative 3’ splice site, A5 = alternative 5’ splice site, MX = mutually exclusive, RI = retained intron, SE = skipped exon.

Integration with Ensembl (GRCh38.113) and Gencode (GRCh38.p14) annotations yielded a combined transcriptome of 620,133 isoforms. Inferring these isoforms with coding sequences and coding potential, we identified 83,409 full-splice matches, referring to transcripts identical to annotated transcripts; 132,108 novel-in-catalog isoforms which use known splice sites in new combinations; and 249,329 novel-not-in-catalog isoforms that contain at least one novel splice site (Figure 1B). Notably, whereas only 4.3% of full-splice match transcripts were predicted to be NMD-sensitive, this fraction was 27% for novel-in-catalog isoforms and 23% for not-in-catalog isoforms. This indicates that NMD-sensitive isoforms more frequently contain novel exons, splice junctions, or novel combinations of exons.

To assess which AS events contribute to NMD-sensitivity of these isoforms, we annotated AS events across the combined transcriptome. Doing so, we identified 1,621,228 AS events that could be divided into five AS-types; 441,957 skipped exons, 218,629 mutually exclusive exons, 292,979 alternative 5’ splice-sites, 316,602 alternative 3’ splice-sites and 351,061 retained intron events (Figure 1C). Next, we quantified the relative abundance of these events using percentage spliced in (PSI) values derived from short-read RNA-seq data for each cell type, under control and NMD-inhibited conditions. This dataset provided the baseline NMD AS events that we subsequently benchmarked using complementary NMD AS prediction approaches.

### 1.2 A splice-context aware method improves NMD AS prediction

To systematically assess the NMD-inducing potential of all AS events in our combined transcriptome, we developed a long-read-based splice-context aware method. This novel method was designed to account for context-dependent NMD events and avoid misattributing NMD-inducing to AS events that co-occur with a true NMD AS events in the same transcript. We compared the performance of this method to the previously applied NMD-exclusive method (Figure 2)(8, 14). This method classifies an AS event as NMD-inducing if it occurs exclusively in isoforms annotated as NMD sensitive, without considering the splice context.

**Figure 2.**
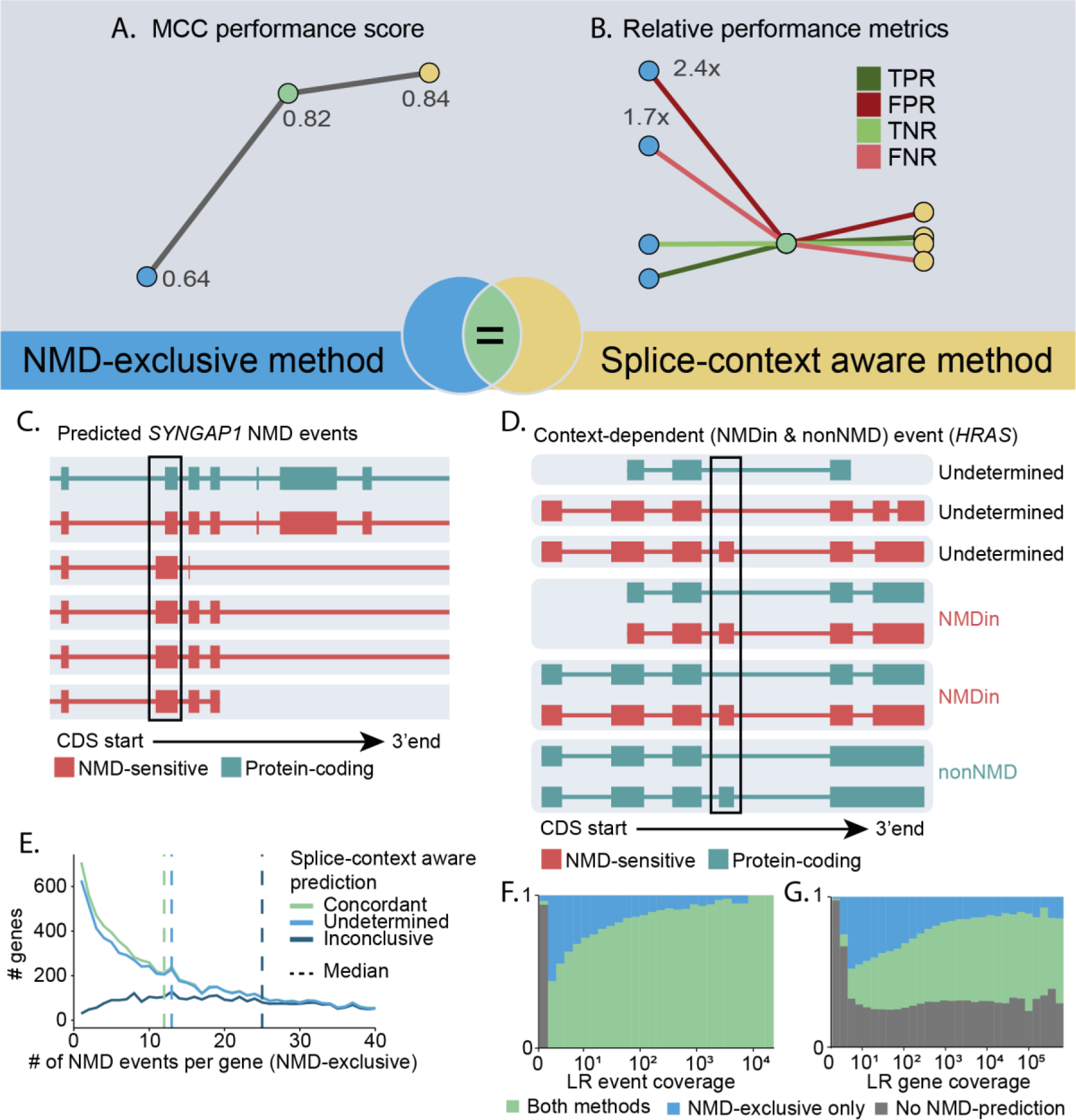
Comparison of two NMD AS prediction methods. **A/B.** Matthews correlation coefficient (MCC) performance scores (**A**) and true positive rate (TPR), false positive rate (FPR), true negative rate (TNR) and false negative rate (FNR) (**B**) for the NMD-exclusive method (blue), splice-context aware method (yellow) and concordant events in the NMD-exclusive method and splice-context aware method (green). **C.** Zoomed-in visualization of a selection of transcripts grouped on splice-context group (highlighted by the light grey background box) in SYNGAP1, a known alternative 3’ splice site NMDin event predicted by the NMD-exclusive method but not the splice-context aware method was highlighted. **D.** Visualization of a selection of HRAS transcripts grouped on splice-context (highlighted by the light grey background box), an AS event that was predicted to be a context dependent NMD AS event by the splice-context aware method. **E.** Number of genes containing 0-40 predicted NMD AS events by the NMD-exclusive method for concordant NMD AS event (green), inconclusive (dark blue) and undetermined (blue) in the splice-aware method. **F/G.** Ratio of AS events receiving a prediction from neither NMD AS prediction methods (grey), both NMD AS prediction methods (green) or only the NMD-exclusive method (blue) over increasing long-read event coverage (minimum long-read coverage for the inclusion or exclusion transcripts) (**F**) and long-read gene coverage (**G**). NMD = nonsense-mediated decay, CDS = coding sequence.

The NMD-exclusive method classified 45% of all AS events, while the splice-context aware method classified a smaller, largely overlapping subset (24%), reflecting its greater stringency (Supplementary Figure 1). AS events labeled only by the NMD-exclusive method tended to have low long-read coverage. The splice-context aware method identified 51% of events supported by fewer than 10 long-read reads, but 90% of events supported by more than 100 reads (Figures 2F, G). This need for higher long-read event coverage by the splice-context aware method also prevented detection of a known NMD AS event in *SYNGAP1*, that was identified using the NMD-exclusive method (Figure 2C).

To compare the accuracy of the NMD-exclusive and splice-context aware method, we evaluated the NMD-inducing potential of predicted NMD and non-NMD events by measuring the ΔPSI upon CHX-mediated NMD inhibition. The splice-context aware method outperformed the NMD-exclusive method (Matthews correlation coefficient (MCC): 0.84 vs. 0.64) and matched the accuracy of concordantly labeled AS events (MCC: 0.82) (Figure 2A). Specifically, the NMD-exclusive method showed a 2.4-fold higher false positive rate and 1.7-fold higher false negative rate (Figure 2B). AS events inconclusive in the splice-context aware method but classified by the NMD-exclusive method showed inflated NMD calls per gene (Figure 2E). This is consistent with the expected misclassification of AS events within genes containing true NMD events.

Additionally, the splice-context aware method, detected 5,481 context-dependent NMD events and 16,249 NMDin & NMDex AS events missed by the NMD-exclusive method (Figure 2D). This includes an NMDin & non-NMD AS event in *HRAS*, whose NMD sensitivity depends on downstream splicing. Within this discordant group, the NMD-exclusive method missed 9.3× more true NMD events, whereas the splice-context aware method produced 1.9× more false positives compared to concordant events (Supplementary Data 2). This indicates that context-dependent AS events can have NMD-inducing potential.

Based on these results, we selected the splice-context aware method for all downstream analyses due to its higher accuracy and ability to detect context-dependent NMD events, despite classifying fewer AS events overall.

### 1.3 Refining candidate NMD AS events by removing technical and biological confounders

Having identified a large set of candidate-NMD AS events, our next step was to refine this dataset into a reliable and biologically meaningful resource. To do so, we systematically assessed both technical and biological parameters that could confound NMD prediction. We first assessed how three technical features influence NMD AS event prediction performance: long-read gene coverage, relative event coverage (supporting event reads / total reads spanning the event region), and the number of splice-context groups supporting each NMD AS event prediction (non-NMD, NMDin, NMDex).

Whereas, excluding AS events with low long-read coverage did not improve performance, removing AS events with low relative event coverage and few supporting splice-context groups did (Figure 3A). Applying a cut-off of >1 splice-context group and >0.1 relative event coverage increased the MCC from 0.84 to 0.90 (Figure 3B). This primarily excluded AS events with low relative event coverage and rare NMDin or NMDex AS events (Figure 3C/D). This effect was not driven by long-read gene coverage (Figure 3E). While excluding AS events with a relative event coverage above 0.1 did not substantially improve NMD AS predictions, AS events with low-to-mid relative event coverage showed atypical PSI distributions. These AS events were characterized by inflated PSI values and greater variability (Figure 3F/G) and mainly caused biased quantification of AS events that share splice junctions with other AS events.

**Figure 3.**
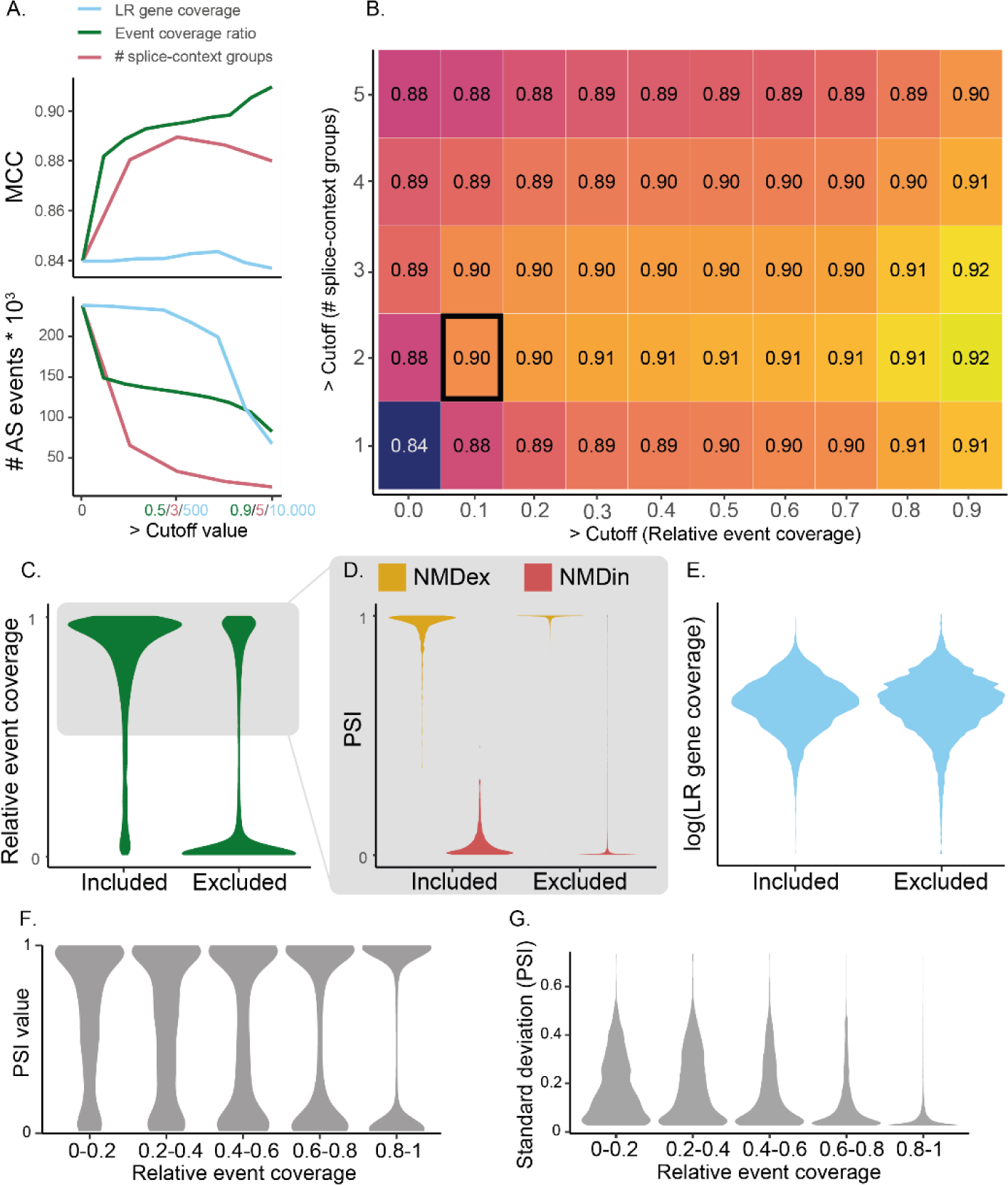
NMD AS prediction parameter cutoff optimization. **A.** Matthews correlation coefficient (MCC) performance and number of included AS events upon increasing cut-off values for long-read gene coverage (blue), relative event coverage (green) and number of splice-context groups (red). **B.** heatmap of MCC scores for different combinations of cut-off values for the number of splice-context groups and relative event coverage. The most optimal combination of cutoff values is highlighted by a black square. **C.** Relative event coverage scores of excluded and included (relative event coverage cutoff >= 0.1, # splice-context groups >= 2) AS events. **D.** PSI of included and excluded events with a relative event coverage of > 50%. **E.** Long-read gene coverage of included and excluded events. **F.** PSI distribution of AS events grouped on relative event coverage. **G.** Standard deviation of the PSI of AS events grouped on relative event coverage. PSI = percentage spliced in, LR = long-read.

After applying the previously identified technical cut-off values, we next addressed biological confounders that can lead to false-positive NMD AS predictions. Retained introns and AS events generating long exons are often NMD-resistant due to nuclear retention, which can result in false identification of NMD AS events (20, 21). Consistent with this, retained intron events did not show increased inclusion of the NMD-causing state in CHX versus DMSO samples, unlike other AS types (Supplementary Figure 2B). Moreover, for NMDin events, the ΔPSI decreased with increasing event length up to 500 bp (R= -0.21, p < 0.0001, Supplementary Figure 2C). Notably, 73% of retained intron NMD-events exceeded 500 bp, compared to 20% for alternative 3’ splice site, 14% for alternative 5’ splice site, and 2% for skipped exon events. The high number of AS events that generate long exons in retained introns, alternative 3’ splice sites and alternative 5’ splice sites explains the NMD-resistance in retained introns and partial resistance observed in alternative 3’ splice site and alternative 5’ splice site events (Supplementary Figure 2D). To avoid these confounding effects, we therefore excluded 8,318 AS events creating exons longer than 500 bp from downstream analyses.

### 1.4 iPSC neural models largely recapitulate NMD-coupled splicing in the human brain

Having validated our NMD AS prediction approach and established cutoff values for identifying high-confidence NMD events, we next asked how this splicing reflects endogenous splicing in the human brain. To address this, we evaluated the presence of supporting reads for the AS events from our experimental data in GTEx bulk short-read RNAseq data across multiple brain regions (Figure 4). For AS events with intermediate PSI (0.1 < PSI < 0.9), up to 69% were supported in GTEx brain region samples, indicating that many of these events are also present in human brain tissue. Support was lower (29%) when including AS events with extreme inclusion or exclusion (PSI < 0.1 or > 0.9). This is likely because these rare splice states require more reads to be reliably detected. Notably, novel AS events were less frequently supported than known events (31% vs. 80%; Figure 4B), reflecting their lower representation in existing transcriptomic datasets.

**Figure 4.**
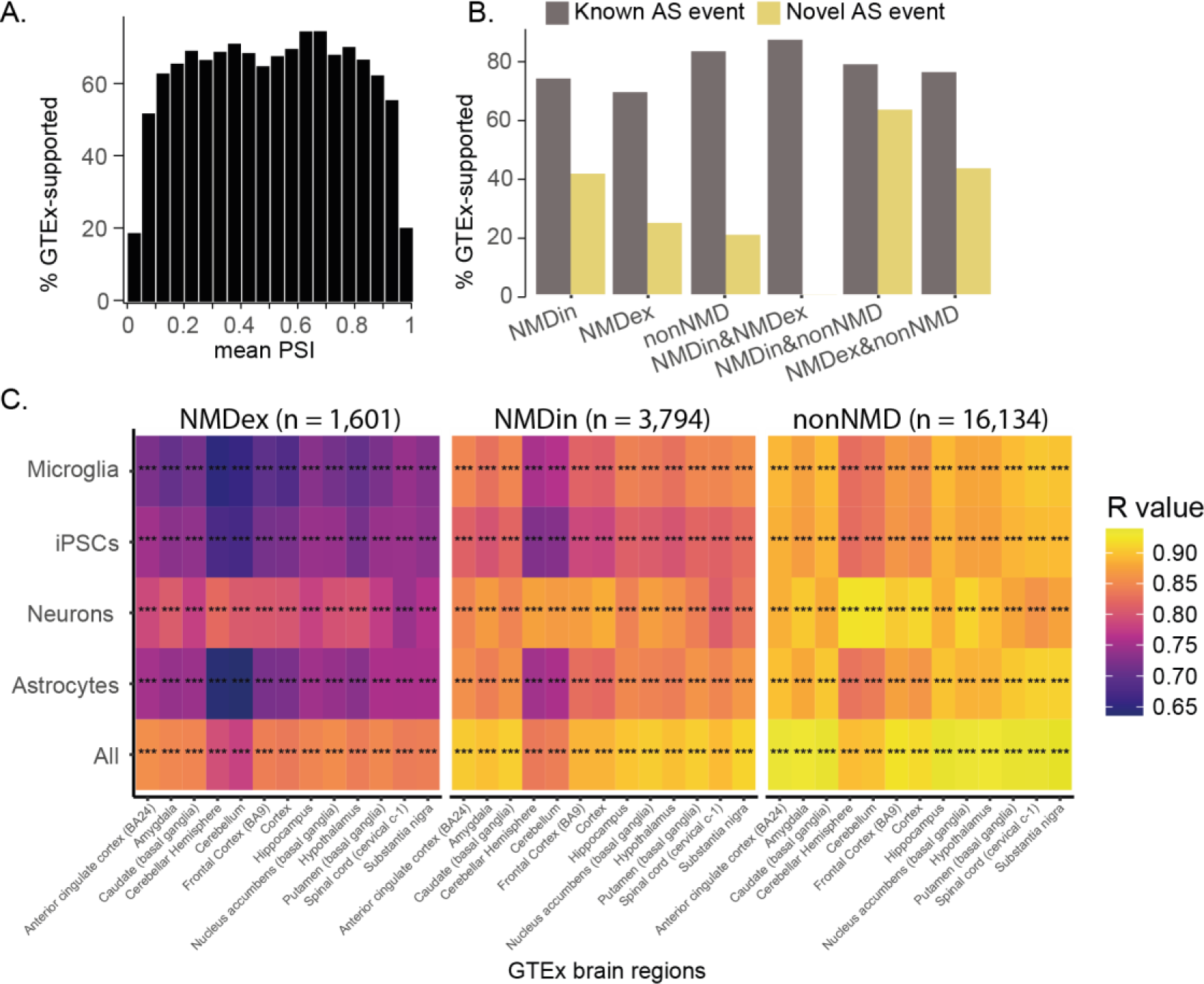
AS event support and PSI correlation in GTEx brain samples. **A.** Proportion of GTEX-supported AS events grouped on their average PSI across all iPSC-derived samples. **B.** Proportion of GTEx-supported AS events for novel and known AS events. **C.** Heatmap of R-values (Pearson correlation) of the average PSI of microglia, iPSCs, neurons, astrocytes and all samples together compared to PSI from GTEx-brain regions per NMD-type. Context-dependent NMD-types were excluded from this analysis. NMD = nonsense-mediated decay, NMDin = AS events predicted to cause NMD upon inclusion, NMDex = AS events predicted to cause NMD upon exclusion, non-NMD = AS events not predicted to cause NMD. * p < 0.05, ** p < 0.01, *** p < 0.001.

To evaluate how well iPSC-derived AS event abundance reflects their abundance in the brain, we compared the PSI of all AS events with intermediate PSI (0.1 < PSI < 0.9) supported in GTEx with the PSI calculated from GTEx bulk short-read RNA-seq across all available brain regions. Overall, mean PSI from our cell models correlated strongly with PSI calculated from the GTEx brain regions (R = 0.63-0.95), indicating that our model broadly reflects endogenous brain splicing (Figure 4C). Correlations were weaker for NMDin (R = 0.74-0.91) and NMDex (R = 0.63-0.87) AS events compared to non-NMD AS events (R = 0.85-0.94), consistent with challenges in quantifying unstable NMD-sensitive transcripts. As expected, averaging PSI across all induced cell types gave the highest correlations scores, consistent with bulk RNA-seq capturing mixed populations. An exception to this was the cerebellum, where neuron PSI correlated most strongly (R = 0.87-0.94 with neurons vs. R = 0.76-0.91 with the multi-cell-type average).

## 2. Characteristics of NMD AS events

To systematically catalog high-confidence NMD AS events, we compiled an online, searchable database of candidate NMD AS events across iPSC-derived brain cell types (https://nadifkasrilab.shinyapps.io/SpliNEx/). We then used this resource to characterize NMD AS event properties.

### 2.1 Novel and known AS events display expected responses to CHX

Across all cell types, the database comprises 15,391 NMDin AS events (26%), 6,750 NMDex AS events (12%), 33,518 non-NMD AS events (57%), and 3,011 context-dependent NMD AS events (5%) (Figure 5A). In total, 40,545 AS events (62%) were novel, defined as AS events where either the inclusion or exclusion isoform was not observed in the reference transcriptome. Novel AS events comprised 79% of NMDin and 78% of NMDex events, substantially higher than the 52% observed in non-NMD events (Figure 5B). Both novel and known NMD AS events, including context-dependent NMD AS events, exhibited CHX-induced ΔPSI changes in the expected directions (ΔPSI > 0 for NMDin, ΔPSI < 0 for NMDex), supporting their NMD-inducing potential (p < 0.0001, Figure 5C, Supplementary Figure 3A, B). Baseline PSI distributions showed that the splice state that triggers NMD (inclusion of the AS event for NMDin events and exclusion for NMDex events) is generally rare, with most transcripts favoring the non-NMD isoform. Specifically, in NMDin AS events (where inclusion triggers NMD) the event was mostly excluded (low PSI), whereas in NMDex AS events (where exclusion triggers NMD) the exon was mostly included (high PSI), relative to non-NMD AS events (Figure 5D). This pattern was even stronger for novel AS events, their NMD-causing splice states were generally rarer, and they showed smaller log ΔPSI shifts upon CHX treatment (Supplementary Figure 3A–D). This may indicate that novel AS events represent lower-abundance or weaker NMD-inducing isoforms.

**Figure 5.**
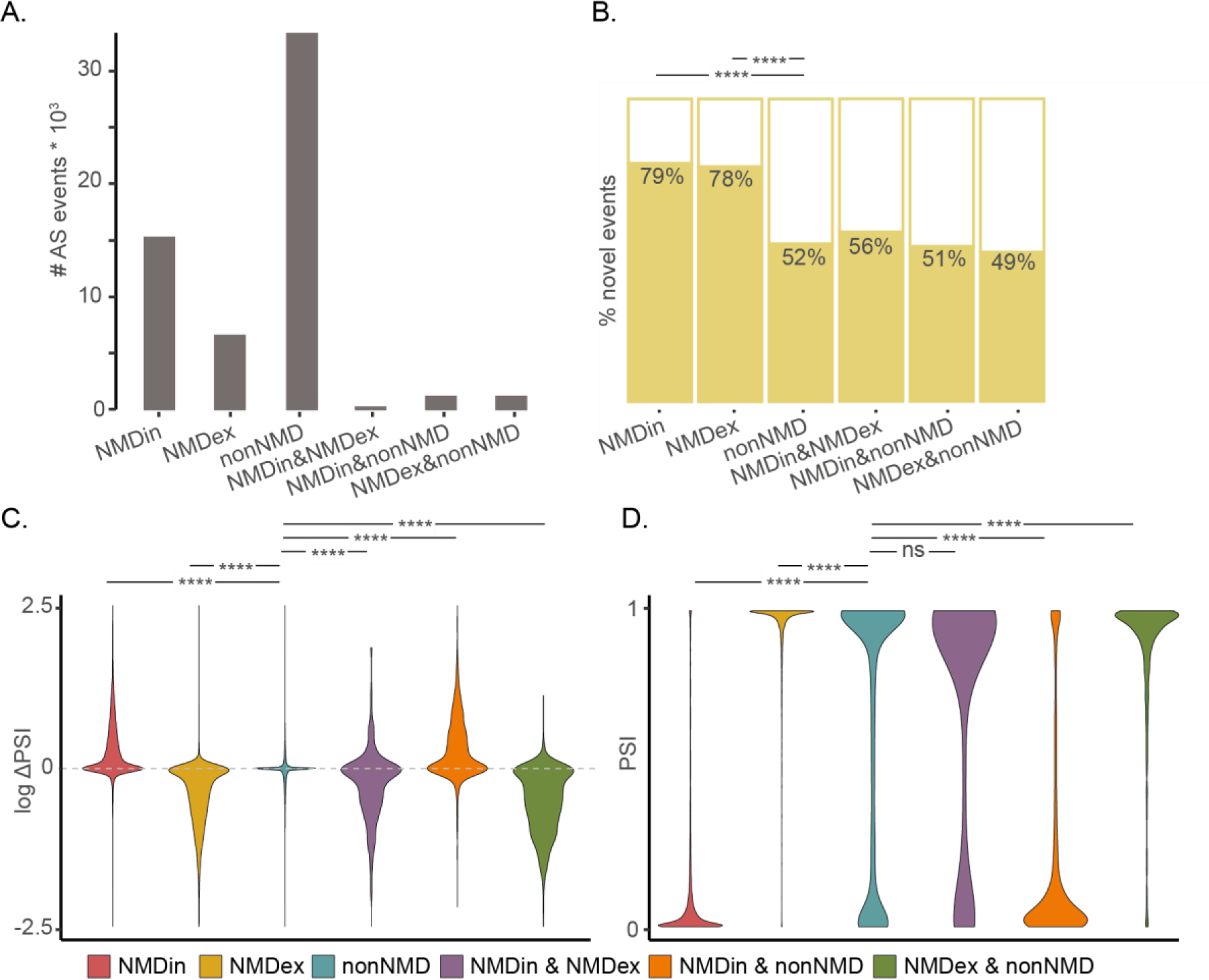
Characteristics of novel and known AS events. **A.** Number of AS events per NMD AS type. **B.**Percentage of novel AS events over the total number of AS events per NMD AS type; significance assessed with Fisher’s exact test. **C.** Violin plots of log ΔPSI (PSI in CHX condition - PSI in DMSO control condition) per NMD AS type; calculated using the two-sided Wilcoxon rank-sum test. **D.** Violin plots of mean CHX-treated PSI value distribution per NMD AS type; calculated using the Wilcoxon rank-sum test. NMD = nonsense-mediated decay, NMDin = AS events predicted to cause NMD upon inclusion, NMDex = AS events predicted to cause NMD upon exclusion, non-NMD = AS events not predicted to cause NMD, PSI = percentage spliced in. NMDin (n = 15,391), NMDex (n = 6,750), nonNMD (n = 33,518), NMDin & NMDex (n = 352), NMDin & nonNMD (n = 1356), NMDex & nonNMD (n = 1303). * p < 0.05, ** p < 0.01, *** p < 0.001 and **** p < 0.0001.

### 2.2 NMD AS occurs in less conserved genes and genomic regions

To gain insight into the biological relevance of NMD AS events, we examined their conservation and explored their potential role in biological processes (using gene ontology terms). Compared to non-NMD AS events with similar PSI, NMD AS events showed generally lower evolutionary conservation at positions across intronic and exonic regions flanking splice junctions, across 100 vertebrate species (Supplementary Figure 4A-E). This pattern was most pronounced for NMDin AS events. For example, skipped exons exhibited a marked reduction in conservation across exonic regions (0.72, fold change [FC] = average conservation in NMD AS events / conservation in non-NMD AS events), exon-intron boundaries (0.71 FC, ±2 nucleotides around the splice site), and flanking intronic sequences (0.76 FC, 0-50 bp from the splice site) but less in more distal intronic sequences (0.83 FC, 50-100 bp from the splice site) (Supplementary Figure 4A-E). In contrast, skipped exon NMDex AS events mainly showed decreases in conservation restricted to intronic regions (0.72 FC, 0-50 bp from the splice site). This includes more distal intronic areas (0.74 FC, 50-100 bp from the splice site) but not in exonic regions (0.99 FC) (Supplementary Figure 4A-E). NMD AS events were strongly depleted in genes under strong selective constraint: only 26% of genes containing NMD AS events had a probability of loss-of-function intolerance (pLI) > 0.8, compared with 41% of genes containing only non-NMD AS events (Supplementary Figure 4F). This suggests that NMD AS events preferentially occur in genes more tolerant to loss-of-function variants.

To determine whether common NMD AS events are found in specific functional gene classes, we analyzed their enrichment across biological processes using gene ontology (GO) terms. Common NMD AS events (>20% average NMD-causing splice state) showed modest enrichment (Observed/Expected number of genes annotated to a given process (OE) between 1 and 2) in biological processes related to cellular component organization (OE = 1.3, p = 0.001), lipid metabolism (OE = 1.9, p = 0.02) and RNA processing (OE = 1.5, p = 0.04) (Supplementary Figure 4G, Supplementary Data 3). Although not statistically significant, the top 50 enriched GO terms also included processes previously linked to NMD AS (2, 22), such as various metabolic processes and RNA splicing (Supplementary Data 3).

## 3. Applications of the NMD AS database: cell-type splicing and therapeutic target identification

To illustrate the potential applications of our NMD AS database, we focus on two use cases: (i) analysis of cell-type-specific differential splicing of NMD and non-NMD AS to investigate AS regulation in fundamental brain development, and (ii) validation, identification, and prioritization of candidate targets for TANGO-mediated upregulation as a therapeutic strategy for NDDs.

### 3.1 Use case I: cell-type-specific splicing patterns of NMD and non-NMD AS events

As a use case, we examined cell-type-specific splicing patterns of NMD and non-NMD AS events to explore the role of AS regulation in fundamental brain development. To this end, we analyzed differential splicing (DS) across iPSC-derived neurons, astrocytes, and microglia. Doing so, we identified 1,817 DS AS events (minimum PSI difference of 0.2) of which 518 were NMD AS events, and 1,299 were non-NMD AS events (Figure 6A). DS AS events were mostly skipped exon events (OR = 1.29, p = 6.2e-12) and NMD AS events were less likely to be differentially spliced compared to non-NMD AS events (OR = 0.74, p = 6.1e-10) (Figure 6B). Notably, DS AS events were enriched in NDD genes (OR = 1.14, p = 0.04), indicating involvement of AS in genes critical for brain development (Figure 6B).

**Figure 6.**
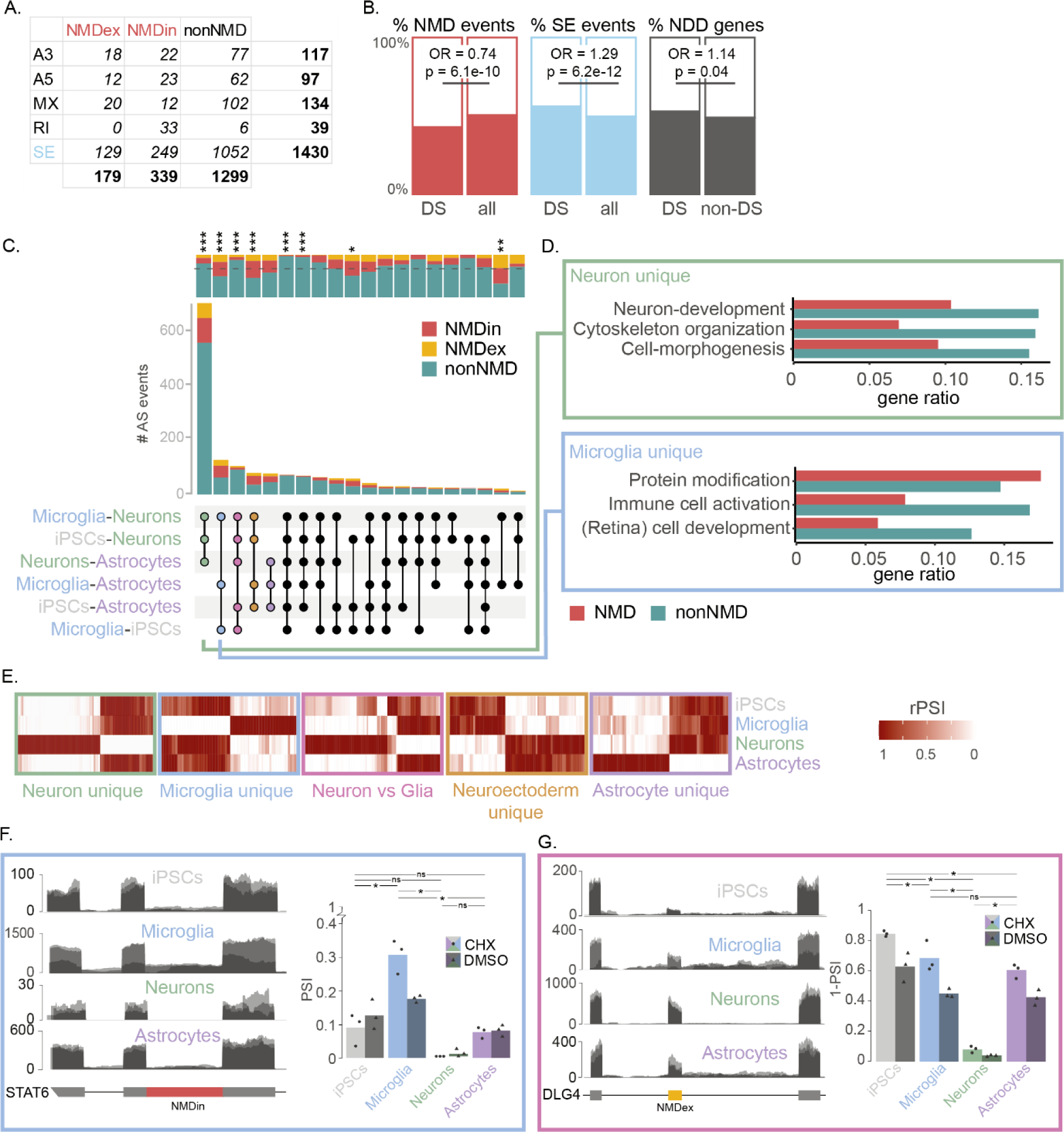
Differential splicing of AS events. **A.** Table of differentially spliced AS events per SUPPA event type (A3, A5, MX, RI, SE) and NMD AS type (non-NMD, NMDin, NMDex). **B.** Ratios of (i) NMD AS events and (ii) skipped exon (SE) events in DS AS events compared to all AS events. (iii) Ratio of NDD genes among genes with and without a DS AS event. **C.** UpSet plot showing combinations of significant pairwise inter-cell-type comparisons (Tukey test) among differentially spliced AS events. Each unique set of comparisons in which an AS event is significant defines an inter-cell-type pattern. Connected dots show the comparisons significant in each pattern, with the bar heights representing the number of AS events showing that pattern. The proportion of NMD versus non-NMD events are annotated above the bars. Enrichment or depletion of NMD events within inter-cell-type splice patterns (indicated by asterisks) was assessed using a two-sided binomial test. **D.** Top three GO-term clusters enriched in neuron-specific and microglia-specific inter-cell-type patterns. Bar height represents the gene ratio of genes containing DS NMD and non-NMD AS events within each GO-term cluster, calculated relative to all NMD or non-NMD DS AS events in the corresponding inter-cell-type pattern. **E.** Heatmaps of relative PSI (rPSI, scaled 0-1 per event, with the lowest PSI set to 0 and the highest to 1) of the top 5 inter-cell-type patterns. The outline colors match the inter-cell-type pattern group colors in panel C. **F/G.** Overlay of short-read RNA-seq coverage from CHX-treated samples and corresponding PSI bar plots across different iPSC-derived cell types (treated ± CHX) for DS AS events in STAT6 and DLG4. Outline colors indicate the inter-cell-type pattern of each event. Statistical significance was calculated using a two-sided Student’s t-test and adjusted for multiple comparisons using the Benjamini Hochberg procedure (*p < 0.05; **p < 0.01; ***p < 0.001; ****p < 0.0001). PSI = percentage spliced in, A3 = alternative 3’ splice site, A5 = alternative 5’ splice junction, MX = mutually exclusive, RI = retained intron, SE = skipped exon, NMD = nonsense-mediated decay, NMDin = AS events predicted to cause NMD upon inclusion, NMDex = AS events predicted to cause NMD upon exclusion, non-NMD = AS events not predicted to cause NMD.

The DS AS events could be divided into 52 distinct inter-cell-type patterns based on which cell-type comparisons showed significant differences, with 18 patterns containing more than 20 AS events (Figure 6C). By far the largest group of DS AS events showed unique splicing in neurons compared to all other cell types and contained 37% of all DS AS events (666/1,817), suggesting that alternative splicing is particularly prevalent in excitatory neurons. Notably, 8 out of 18 of the inter-cell-type patterns with more than 20 AS events showed significant enrichment or depletion of NMD AS events. For example, NMD AS events were enriched in microglia-unique and astrocyte-unique splicing clusters, constituting 49% and 46% of the DS AS events, respectively (Figure 6C). Among the AS events in the top inter-cell-type splice patterns was an NMDex AS event in *DLG4*. This NMDex AS event was primarily included in mature neurons (96%) and therefore not degraded in neurons, consistent with prior reports (Figure 6G) (4). Besides that, we also identified other NMD AS events that were not previously known to be differentially spliced, including in *EZH2, STAT6, P2RX4, KANSL2* and *CHD2* (Figure 6F, Supplementary Data 4).

To explore cellular functions associated with DS AS events, we performed gene set enrichment analysis on genes harboring DS AS events within the top five inter-cell-type patterns. Although statistical power was sometimes limited by the number of DS AS events, clustering of the top 100 GO-terms revealed clear cell-type-specific trends that reflect known biology (Figure 6D, Supplementary Data 5). For example, neuron-unique splicing was linked to neuronal development, microglia-unique splicing to immune activation and immune-cell development, and neuron vs glia splicing to cell motility, a function shared by microglia and astrocytes (Figure 6D, Supplementary Data 5). DS non-NMD AS events generally contributed most to the enrichment of these clusters, with 1 - 2.4-fold higher gene ratios than NMD AS events across enriched terms for neuron-specific splicing. Notable exceptions included a GO-term cluster related to post-translational modification in the microglia-unique pattern, where NMD AS events contributed slightly more than non-NMD AS events (1.2-fold), as well as GO-term clusters related to blood vessel regulation (1.1-fold) and metabolism & protein modification (1.4-fold) in the astrocyte-unique inter-cell-type pattern (Figure 6D). Although these differences are not statistically significant, these trends suggest that DS NMD AS events may contribute more to some cell type-specific biological processes than to others.

#### 3.1.1 Regulation of differentially spliced AS events varies between brain cell types

To understand how NMD-inducing AS is differentially regulated across cell types, we clustered AS events within the previously identified inter-cell-type splicing patterns into groups of co-spliced AS events based on their inclusion and exclusion profiles between cell-types. Interestingly, for 4 of the top 5 inter-cell-type groups NMDin and NMDex AS events segregated into mirrored clusters, with NMD-causing isoforms predominating in specific cell types (Supplementary Figure 5A). For instance, microglia-specific DS AS events consisted of two clusters: one with DS AS events that show high inclusion in microglia (including 41 out of 42 NMDin events) and another cluster with DS AS events that show high exclusion in microglia (including 21 out of 22 NMDex events). These mirrored patterns suggest that DS-NMD AS events may be regulated in a manner that is specific to the NMD AS event type.

To investigate regulators of cell-type-specific NMD AS, we focused on PTBP2, a neuronally enriched splicing factor critical for controlling AS, including NMD AS events, during neuronal differentiation (7, 23). Interestingly, skipped exon events in cluster 1 of the neuron unique inter-cell-type pattern (p = 2.3*10-4, OR = 4.5, n = 399) and cluster 1 of the neuron vs glia specific inter-cell-type pattern (p = 0.04, OR = 10.44, n = 60) were significantly enriched in DS AS events that were differentially spliced upon knockdown of PTBP2 (Supplementary Figure 5B). In both cases, this enrichment was specific to the clusters containing mostly NMDin AS events, except for an NMDex AS event and known target of PTBP2 in *DLG4* (Supplementary Figure 5A/B). Interestingly, all DS AS events that were also differentially expressed upon *PTBP2* knock down in the neuron-specific and neuron vs glia NMDin-enriched clusters showed increased inclusion upon *PTBP2* knockdown, indicating that PTBP2 normally represses the inclusion of these DS AS events (Supplementary Figure 5C).

By repressing the inclusion of NMDin events (which trigger NMD when included) PTBP2 may maintain RNA levels of these genes. Indeed, genes containing DS NMD AS events in cluster 1 of neuron-unique events were 5-fold more often downregulated than upregulated upon *PTBP2* knockdown. On the other hand genes containing only DS non-NMD AS events showed a more balanced distribution (1.3-fold down vs. upregulated) (Supplementary Figure 5D). These results indicate that PTBP2 may influence gene expression by modulating inclusion of NMDin AS events, highlighting RBPs as potential mediators of cell-type-specific post-transcriptional regulation through NMD AS.

### 3.2 Use case II: Identifying abundant NMD AS events in autosomal-dominant NDD genes as potential therapeutic targets

As a second use case, we aimed to identify NMD AS events that could be therapeutically targeted to upregulate wild-type allele protein levels in NDDs caused by heterozygous loss-of-function variants. To this end, we systematically analyzed our NMD AS dataset for events occurring in reported AD NDD genes (9). This analysis identified 936 NMD AS events in 250 AD NDD genes (63% of all 394 AD NDD genes with AS event annotations in our dataset) as potential TANGO targets (Figure 5A). Notably, an NMDin AS event in *SCN1A*, the only gene for which TANGO has been tentatively successfully applied in clinical trials (11), showed one of the highest abundance of any NMD AS event in an AD NDD gene, with 81% inclusion in neurons. Since TANGO efficacy likely depends on targeting abundant NMD AS events, many of the 936 NMD AS events identified may not be suitable candidates due to low occurrence of the splice state that causes NMD (inclusion of the AS event for NMDin events and exclusion for NMDex events). By selecting only NMD AS events with more than 10% NMD-causing splice state (PSI > 0.1 for NMDin; PSI < 0.9 for NMDex) in at least one cell type, we further refined our focus to 145 abundant NMD AS events in 81 AD genes as potential high-frequency TANGO targets (Figure 7A, Supplementary Data 6).

**Figure 7.**
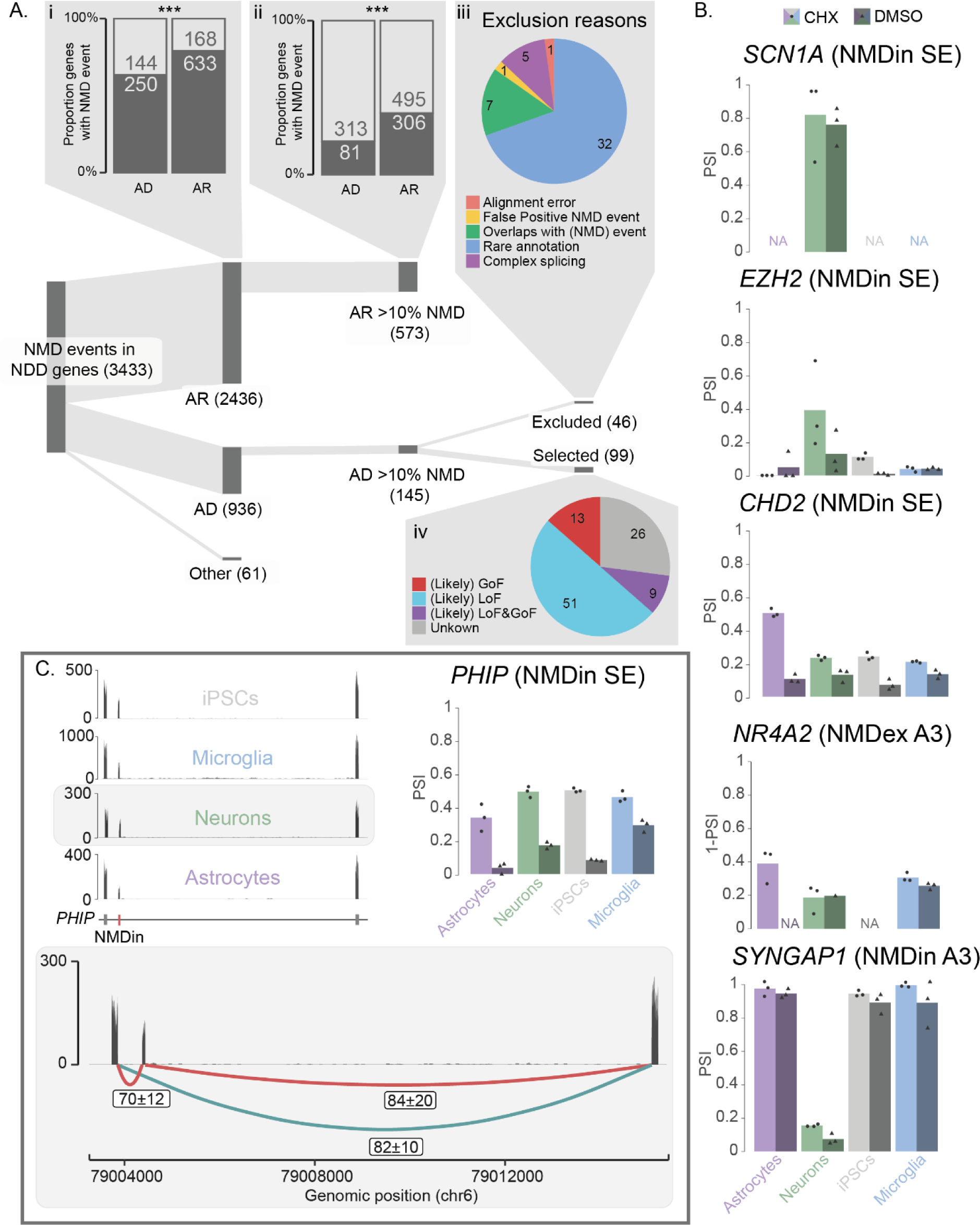
Identification of NMD AS events in NDD genes. **A.** Sankey diagram of NMD AS events in NDD genes. Bar plots show the proportion of NDD genes related to autosomal dominant (AD) and autosomal recessive (AR) disorders containing a NMD AS event (i) or a common (>10%) NMD AS event (ii). The pie charts indicate reasons for exclusion after manual assessment and the molecular mechanism of disease for NDD-associated genes containing an NMD AS event (selected after manual assessment). **B.** PSI of previously reported NMD AS events across different iPSC-derived cell types, with and without cycloheximide (CHX) treatment. **C.** Coverage plots of the CHX treated condition per cell-type with replicates overlaid (upper left) and PSI of novel NMD AS events across different iPSC-derived cell types, with and without CHX treatment (upper right) with zoom in on coverage and splice-junction reads of neuron CHX treated condition (bottom). PSI = percentage spliced in, NMD = nonsense mediated decay, NDD = neurodevelopmental disorders. LoF = loss-of-function, GoF = gain-of-function, NMDin = AS events predicted to cause NMD upon inclusion, NMDex = AS events predicted to cause NMD upon exclusion, non-NMD = AS events not predicted to cause NMD.

Because selecting for abundant AS events can enrich for AS events with inflated PSI values due to low-to-medium relative event coverage (Figure 3F/G), we manually assessed read coverage and checked for interfering splicing in the surrounding region. We excluded 46 NMD AS events, of which 32 were removed due to unreliable PSI estimates arising from spurious AS event annotations, 7 due to unreliable PSI estimates arising from overlapping AS events and 5 due to unreliable PSI estimates because of complex splicing in the region (Figure 7A, Supplementary Data 6). These NMD AS events had low-to-medium relative event coverage scores which artificially inflated their NMD-causing splice state to above the 10% threshold. Two additional AS events were excluded because of an alignment error (n = 1) or false positive NMD AS event (n = 1) (Figure 7A).

The 99 remaining NMD AS events were associated with 69 NDDs; of these, 60 AS events involving 42 NDD genes which are known to have haploinsufficiency as their disease mechanisms (likely loss-of-function or likely loss-of-function & gain-of-function), making them candidates for TANGO (Figure 7A, Supplementary Data 6). For 26 other AS events in 16 genes, the molecular mechanism of disease is still unresolved but may also represent potential TANGO candidates (Figure 7A, Supplementary Data 6). An additional 13 AS events were found in genes associated with a gain-of-function molecular mechanisms (Figure 7A, Supplementary Data 6). Potential TANGO targets included a known abundant NMDin AS event in *SCN1A* (PSI 81%) and NMDex AS event *DLG4* (percentage spliced out (PSO, defined as 1-PSI) 8%-85% across cell types), as well as previously reported NMDin AS events in *CHD2* (PSI 21%-49%) and *EZH2* (PSI 5%-39%), and an NMDex AS event in *NR4A2* (PSO 19%-39%) (Figure 7B, Supplementary Data 6). We also identified novel candidate targets with a high NMD-inducing splice state, including an NMDin AS event in *PHIP* (PSI 33%-49%), in which AD loss-of-function variants cause Chung-Jansen syndrome (OMIM: 617991) (Figure 7C/D, Supplementary Data 6). This event showed a significant difference in PSI between DMSO (PSI 4%-29%) and CHX (PSI 33%-49%) in all cell types (p = 0.014-0.004), supporting that its inclusion is targeted by NMD.

## Discussion

The successful application of the AON-based TANGO strategy for Dravet syndrome in clinical trials provides hope for patients and families affected by NDDs. While previous studies have reported a wealth of NMD-sensitive AS events, including in NDD-related genes (8, 14, 15), information on their presence and occurrence in disease-relevant cell types is limited. Here, we leveraged long- and short-read transcriptome sequencing in iPSC-derived brain cells and developed a more accurate splice-context-aware method to identify NMD AS events. To facilitate further research into non-NMD and NMD AS and the therapeutic potential of NMD AS events, we generated a comprehensive database of NMD and non-NMD AS events and provide a user-friendly web tool to explore the presence and abundance of AS events (https://nadifkasrilab.shinyapps.io/SpliNEx/). The interface includes clear, accessible descriptions of key terms and concepts needed for navigation and interpretation of the dataset.

To demonstrate the utility of our resource, we present two use cases. In one use case, we applied our resource to identify NDD-associated genes with abundant NMD AS events. We identified known NMD AS events in genes such as *SCN1A*, *DLG4*, *CHD2*, *NR4A2*, and *EZH2* (*2, 4, 8, 10*), and uncovered novel abundant NMD AS events, including an NMD AS event in *PHIP*. In contrast, many previously reported NMD AS events were deemed unlikely to be effective TANGO targets in our analysis, as they either failed stringent quality cutoffs, were not predicted to be NMD-sensitive, or were not abundant in brain cells (Supplementary Figure 6A)(8). This suggests that TANGO’s applicability may be more limited than initially anticipated. Indeed, expanding on previous systematic analyses while focusing on abundant NMD AS events, highlights 12 NDDs as the most promising candidates for TANGO, compared to 53 previously identified (Supplementary Figure 6A)(9). Nevertheless, upregulation in supportive tissues or cell types with higher levels of relevant NMD AS events, not included in this study, could provide therapeutic benefit. Extensive *in vitro* and *in vivo* studies will be required for each NDD and candidate NMD AS event to determine how much gene upregulation can be achieved, whether it restores normal expression, and if it can reverse or halt the associated phenotype.

Beyond confirming the presence of potentially targetable NMD AS events in autosomal dominant neurological disorders, our resource can also be used to identify and prioritize both novel and known NMD AS events for therapeutic exploration in other contexts. For example, in autosomal-recessive disorders with residual enzyme activity, TANGO could upregulate partially active proteins, as demonstrated *in vitro* for *PCCA* (13). Our dataset offers insight into the presence and abundance of NMD AS events in these genes. Additionally, our resource can help identify NMD AS events in genes that could be upregulated to modulate disease-relevant pathways, even when the genes are not directly causative. For instance, we identified a highly abundant NMDin AS event in *RANBP17*, with up to 65% nonproductive splicing in neurons, whose upregulation may be leveraged to treat early-onset isolated dystonia (24). Similarly, in mice, overexpression of *Nr4a2* using AAVs has been shown to reduce inflammation and promote neurogenesis in brain injury and neurodegenerative disorders unrelated to genetic variants in *NR4A2* (25–27). TANGO might provide a means to achieve this upregulation in patients. Finally, our methods can be applied to long-read and short-read datasets from other cell types of interest for TANGO, such as retinal cells for inherited eye-related disorders. However, cut-off values for relative event coverage and the number of splice-context groups may need to be adjusted depending on dataset depth.

We show that our resource can be leveraged to uncover brain splicing programs and gain insight into the regulatory mechanisms that shape them. Consistent with previous studies highlighting AS as a key regulator of neuronal differentiation and function (28, 29), most DS AS events showed neuron-specific splicing and included a subset of AS events likely regulated by PTBP2. At the same time, our resource may help identify and explore lesser-known groups of DS AS events. For example, we identified a substantial set of DS AS events that are microglia-specific, representing a potentially important and underexplored splicing program that may be further investigated using our resource. This group includes a retained intron in *STAT6*, a key regulator of microglial immune response (30). This raises the possibility that differential intron retention, known to modulate immune activation in macrophages (21), may play similar roles in microglia. Furthermore, DS AS events with unique splicing in microglia were enriched in NMD AS events and formed mirrored clusters of NMDin and NMDex AS events, reminiscent of PTBP2-mediated regulation in neurons. Investigating whether specific splice regulators underlie this patterns may provide insights into brain splicing programs and can and guide the rational design of AONs for therapeutic applications (7).

Both NMD-inducing potential predictions and AS quantification should be interpreted with caution. Manual review of abundant events in autosomal dominant NDD genes showed that especially low-to-medium relative event coverage can inflate PSI estimates. Because we prioritized abundant NMD AS events, such cases with inflated PSI values are overrepresented and do not reflect the global false-positive rate. Nevertheless, event-coverage-dependent PSI inflation should be considered for all application of our resource, particularly when focusing on abundant NMD AS events. To support users, the webtool highlights events with low relative event coverage.

Our methods also have scope limits. The splice-context-aware approach requires sufficient long-read coverage for both isoforms (representing the inclusion and exclusion). Hence, rare AS events or AS events in poorly covered genes may go undetected by this method (e.g. the NMD-causing event in SYNGAP1). We also do not capture NMD AS events outside coding sequences that may create alternative start sites. Finally, some biological factors such as NMD resistance in long exons or context-dependent splicing may prevent degradation of transcripts with predicted NMD-inducing AS events (20). In all these cases, evaluating the unfiltered dataset, together with PSI comparisons between CHX-treated and control samples, can provide complementary evidence for NMD sensitivity. To aid evaluation, the webtool integrates splice-junction context and links to the full long-read transcriptome in the UCSC Genome Browser.

Splicing in our iPSC-derived cells reflect cellular composition across brain regions: average PSI across all cell types mirrors each region’s mixed cell-type makeup, while neuron-specific PSI aligns most closely with cerebellar splicing, consistent with the cerebellum’s high neuronal content (31). Support for novel AS events in human brain data was less consistent, likely reflecting differences between iPSC-based models and the physiological context of human brain tissue. However, these differences may also arise from the rarity of novel AS events or their restriction to specific cell populations, which can be diluted in bulk tissue GTEx data, underscoring the value of our cell type-resolved resource. Importantly, our dataset and observations represent a snapshot influenced by genetic background, cellular context, developmental stage and the environment. AS events warrant validation across contexts and donors. In stimulus-responsive microglia and astrocytes, immune activation may shift (NMD) AS abundance (32, 33).

In summary, we provide a user-friendly resource to evaluate (NMD) AS, prioritize TANGO targets for NDDs and other disorders, and generate hypotheses about splicing regulation in the brain. Our detection and quantification framework is portable to other datasets, tissues, and cell types. Future work should test AS across additional contexts and genetic backgrounds and assess the functional and clinical targetability of NMD-associated events, with the goal of advancing TANGO-based therapies for monogenic NDDs.

## Methods

### iPSC differentiation into neurons, astrocytes and microglia

All induced cell types in this study were derived from the control line UCSFi001-A (Coriell Institute; GM25256, RRID: CVCL_Y803), reprogrammed from skin fibroblasts of a healthy 30-year-old male. IPSC culture and maintenance were performed according to previously established protocols used for all downstream differentiations (16), including those for neurons, astrocytes, and microglia. Briefly, iPSCs were maintained on Geltrex-coated plates in Essential 8™ Flex Basal-based medium at and routinely passaged when they reached approximately 80–90% confluence. To generate glutamatergic neurons, a doxycycline-inducible *Ngn2* gene cassette was introduced into iPSCs using a previously described protocol (16, 17). For the iPSC condition, iPSCs, harboring the inducible *Ngn2* cassette, were plated and maintained as earlier described in the absence of doxycycline to preserve their pluripotent state (16). For the neuron condition, iPSCs, harboring the inducible *Ngn2* cassette, were plated and cultured in the presence of doxycycline to generate cortical glutamatergic neuron following an established protocol (16). Rat-astrocytes were added to support maturation of the culture at days *in vitro* (DIV) 2. For iPSC-derived astrocytes and microglia, iPSCs were differentiated using previously established methods (18, 19). In short, astrocytes were differentiated from iPSCs lacking the Ngn2 cassette using astrocyte Medium (ScienCell®), while microglia were generated from the same iPSCs via stepwise differentiation through hematopoietic progenitors and subsequent maturation in microglia differentiation medium as previously described (18, 19).

### Optimization of CHX concentration and treatment duration for NMD inhibition

To estimate the optimal concentration and duration of CHX treatment for NMD inhibition, a timeline and dosage test was performed on neurons at DIV21. The efficacy of NMD inhibition was assessed by measuring inclusion of an NMD AS event in *CHD2* using RT-PCR. Neurons were treated with 0, 20, 50, or 100 µg/mL CHX for 5, 8, or 16 hours. Following treatment, cells were harvested in DNA/RNA Shield (Zymo Research, #R1200-125), and total RNA was extracted using the Quick-RNA Microprep Kit (Zymo Research, #R1055) according to the manufacturer’s instructions. cDNA synthesis was performed using the iScript cDNA Synthesis Kit (Bio-Rad, #1708890). RT-PCR reactions were carried out using AmpliTaq 360 DNA Polymerase mix (applied biosystems, #4398886) with primers flanking a previously reported NMD AS event in *CHD2* (Forward: 5’-GGTGGAAGATGATTCTCGCC-3’, Reverse: 5’-CAACAAGTAATCCGCTCGGG-3’). PCR products were resolved on a 1.5% agarose gel, and the extent of NMD inhibition was evaluated by quantifying the ratio of NMD-sensitive (bands at 192 + 181 bp) to total band intensities (bands at 192 + 181 + 169 bp) using GelAnalyzer. This revealed consistent and comparable NMD inhibition across all tested CHX concentrations and treatment durations (Supplementary Figure 7). To confirm that different cell types respond similarly to CHX, astrocytes (DIV70), microglia (DIV23), and iPSCs (48h upon plating as single cell) were treated with CHX concentration ranging between 10-100 µg/mL for 5 hours, after which NMD-inhibiting potential was evaluated. Since, all cell types showed similar NMD-inhibition between CHX concentrations (Supplementary Figure 7), a final treatment condition of 20 µg/mL CHX for 5 hours was selected for all cell types for RNA-seq sample generation.

### Long-read and short-read RNA-sequencing sample generation

To obtain RNA from NMD-inhibited and vehicle brain cell types for long-read and short-read RNA sequencing, cells were plated according to previously established protocols as follows: 6 wells × 300,000 cells for iPSCs (16), neurons (16), and astrocytes (18) in 6-well plates, and two batches of 12 wells × 250,000 cells for microglia (19). Cells were treated with 20 µg/mL CHX or an equivalent volume of DMSO for 5 hours: neurons at DIV49, astrocytes at DIV41, microglia at DIV23, and iPSCs 48 hours after plating. After 5 hours of CHX treatment, all samples were harvested in DNA/RNA Shield (Zymo Research, #R1200-125) and RNA was isolated using the Quick-RNA Microprep kit (Zymo Research, #R1055) according to the manufacturer’s protocol. For microglia, four wells (two from each batch) that received the same treatment were pooled to create one single replicate in order to minimize batch variability. All together this resulted in three samples treated with CHX and three samples treated with DMSO for each cell type (iPSCs, neurons, microglia and astrocytes). After RNA isolation the RNA quality was assessed on the Agilent’s Tapestation system, yielding RNA integrity number (RIN) values of 6.9-9.1 for neurons, 9.0-9.6 for iPSCs, 9.4-9.8 for astrocytes, and 8.4-9.6 for microglia.

### Short-read RNA sequencing and data processing

For short-read RNA sequencing mRNA was enriched using mRNA capture. IPSC and neuron samples were sequenced on 2×75 PE cycle flow cells Nextseq 550 and astrocyte and microglia samples were sequenced on a 2×100bp PE cycle flow cells using NextSeq2000. To compensate for the loss of coverage on the human genome due to the co-culture of rat-astrocytes of the neurons, twice the amount of neuron samples were loaded on the flow cells. This resulted in the average number of reads ranging between 83M-122M in neurons, 48M-69M in iPSCs and 52M-88M reads for astrocytes and microglia. Data quality was assessed using FASTQ. For neurons samples, reads mapping to the rat genome (originating from the co-cultured rat astrocytes) were removed using BBMap Seal (v39.01) with human (hg38) and rat (Rnor 6.0) reference genomes and the option ambig=toss. For iPSCs, astrocytes, and microglia, all reads were aligned to the hg38 reference genome using STAR (v2.7.10b) (34), whereas for neurons only the remaining human reads were aligned after removal of rat-derived reads.

### Long-read RNA sequencing and data processing

For all cell types, two samples with RNA Integrity Numbers (RIN) greater than 7.0 (one treated with CHX and one DMSO-control) were selected for long-read sequencing using PacBio technology (Iso-Seq). Libraries for long-read sequencing were prepared using the SMRTbell Prep Kit 3.0 according to the manufacturer’s instructions, with enrichment for transcripts longer than 3 kb. All samples passed the quality control steps outlined in the manufacturer’s protocol. IPSCs and neurons were sequenced on the PacBio Sequel II system and astrocytes and microglia were sequenced on the PacBio Revio system, using one 8M SMRT cell per sample. Raw movie files were processed using PacBio CCS (v4.0) to generate circular consensus sequences (HiFi reads) on the Sequel II or Revio system. Barcodes were removed using LIMA (v.2.7.1) and trailing poly(A) and poly(T) tails were clipped using the isoseq3 refine tool (v.4.0.0). For neuron samples, reads mapping to the rat genome (originating from the co-cultured rat astrocytes) were removed using BBMap Seal (v39.01) with human (hg38) and rat (Rnor 6.0) reference genomes and the option ambig=toss. All (human) full-length reads were then aligned to the hg38 reference genome using pbmm2 (v.1.12.0), the PacBio implementation of minimap2. TAMA cluster and TAMA merge were then used to construct one combined transcriptome by integrating reads from all samples (35), along with transcript annotations from Ensembl (GRCh38.113) and NCBI (GRCh38.p14) while maintaining source information for each transcript. The coding potential of each transcript in the assembled transcriptome was then predicted using GeneMarkS-T (v.5.1) (36), included in SQANTI3 (v5.1.1) (37). To incorporate short-read evidence for AS events, splice junction (SJ) files obtained from the short-read alignment generated by STAR (v2.7.10b) were provided as input for SQANTI3 (34). Finally, SUPPA2 (v2.3) was used to identify local AS events (38). Alternative splicing event types (A3, A5, MX, and SE) detected by SUPPA2 were quantified using the percentage spliced in (PSI), defined as the ratio of inclusion reads to the sum of inclusion and exclusion reads and reflecting the relative usage of a specific AS event, as previously described (8). Retained intron (RI) events were quantified separately using rMATS (39).

### NMD AS event prediction

Two approaches were used to identify NMD AS events; an NMD-exclusive method and an splice-context aware method. Although AS in the 5’UTR or 3’UTR might trigger NMD, the prediction of both methods were based on only transcripts with splicing events occurring within their predicted coding sequences. The NMD-exclusive method, labels events as NMD if they occur exclusively in transcripts predicted to undergo NMD by GeneMarkS-T (Supplementary Figure 8). In non-NMD events, both inclusion and exclusion isoforms were predicted to be protein-coding. In NMDin events, all inclusion isoforms were predicted to undergo NMD, while exclusion isoforms were protein-coding. Conversely, in NMDex events, exclusion isoforms were NMD-sensitive, and inclusion isoforms were protein-coding. The splice-context-aware method grouped all identified transcripts sharing the same intron chain, strand, and coding sequences start site. Within each splice-context group, events were classified as follows (Supplementary Figure 8):

- non-NMD: all isoforms supporting inclusion or exclusion of the splicing event were predicted to be protein-coding.
- NMDin: all isoforms supporting inclusion of the AS event were predicted to be NMD-sensitive, while those supporting exclusion were predicted to be protein-coding.
- NMDex: all isoforms supporting exclusion of the AS event were NMD-sensitive, while those supporting inclusion were protein-coding.
- Inconclusive: both isoforms supporting the inclusion and exclusion of the AS event were predicted to be NMD-sensitive and NMD AS prediction was therefore inconclusive.
- Undetermined: isoforms within the splice-context group only supported inclusion or exclusion of the event and could therefore not be assessed.

A consensus NMD status was then determined for each event based on whether any splice-context groups supported non-NMD, NMDin, or NMDex. If multiple splice-context groups for a single AS event supported conflicting classifications (e.g., both NMDin and non-NMD), the AS event was labeled as context-dependent, using combinations such as NMDin & non-NMD, NMDex & non-NMD, or NMDin & NMDex.

### NMD AS event prediction validation

To evaluate the performance of the NMD AS prediction methods, we assessed whether predicted AS events showed significant log-fold PSI (ΔPSI) changes in the expected direction upon CHX treatment using a two-sided paired t-test across all pooled samples. NMDin events were expected to show positive ΔPSI (higher PSI in CHX-treated samples), while NMDex events were expected to show a negative ΔPSI (lower PSI in CHX-treated samples) upon CHX treatment. No correction for multiple testing was applied, as the goal was not to identify individual significantly regulated AS events but to assess the overall predictive performance of the method. AS events were classified as follows:

- True positives: NMD AS events with a significant log-fold PSI change in the expected direction upon CHX treatment (i.e., increased PSI for NMDin events or decreased PSI for NMDex events)
- True negatives: predicted non-NMD events that showed no significant log-fold PSI change
- False positives: predicted NMD-targeted events with a significant log-fold PSI change in the opposite direction (i.e., decreased PSI for NMDin events or increased PSI for NMDex events)
- False negatives: predicted non-NMD events that showed a significant log-fold PSI change upon CHX treatment

Based on these classifications, we calculated standard performance metrics including true positive rate (TPR), true negative rate (TNR), false positive rate (FPR), false negative rate (FNR) and Matthews correlation coefficient (MCC) to quantitatively assess prediction accuracy. MCC was chosen because it provides a balanced measure of performance that remains robust to class imbalance, avoiding bias from inflated numbers of true and false negatives due to limited statistical power. The MCC score and other performance metrics were calculated for the NMD-exclusive and splice-context-aware methods to assess method performance.

### Optimizing Parameters for NMD AS prediction

After selecting the splice-context aware method as preferred method for NMD AS prediction due to its higher accuracy, we evaluated the impact of three parameters: the number of splice-context groups supporting an NMD AS prediction, long-read gene coverage (total long-read reads mapping to a gene) and relative event coverage. Relative event coverage was defined as the proportion of AS event-supporting reads (transcripts supporting the inclusion or exclusion of the AS-event) relative to the total long-read region coverage, reflecting the contribution of the event coverage to the long-read gene coverage that can be used to estimate the relevance of an AS event annotation. First, MCC and other performance metrics were calculated across cut-offs for each parameter individually: long-read gene coverage (0, 10, 50, 100, 500, 1000, 5000, 10,000 long-read reads), relative event coverage (≥0-90% in 10% increments), and the number of splice-context groups (≥1-5). As long-read coverage showed no effect on performance, it was excluded from further analyses. MCC and other metrics were then evaluated across all cut-off combinations of relative event coverage and splice-context group number. Based on these results, an optimal threshold of ≥2 supporting splice-context groups and ≥10% relative event coverage was identified. To further assess biological and technical AS event properties affecting NMD-sensitivity and PSI quantification reliability, the relationship between exon length and NMD inhibition was assessed by correlating exon length (bp) with the log ΔPSI. To remove NMD-resistant AS events, AS events leading to an exon length > 500 bp were excluded. In addition, the effect of short-read junction coverage on PSI estimation was evaluated by analyzing PSI distributions across total short-read junction coverage from three replicates per condition. To avoid artificially simplified PSI distributions, PSI values derived from junction coverages <30 were excluded. Altogether, applying the established cut-offs created a final data set of 58,670 AS events.

### Comparing AS in iPSCs with brain region GTEX data

To assess how splicing in our database reflects endogenous splicing in the human brain we obtained SJ files (STAR) from GTEx brain region samples (v10). For each AS event, the number of GTEx reads covering the splice junctions supporting the AS events in our database was calculated. GTEx supported AS events were defined as AS events where all related splice junctions showed at least 1 splice junction read in any of the samples. For all A3, A5, MX, and skipped exon events that were supported in GTEx, the PSI was calculated using the earlier described method. Retained intron events were excluded from the analysis. For the PSI correlation analysis over the different GTEx brain regions, context-dependent NMD AS events were excluded.

### Creating an interactive database for NMD AS event exploration

To provide accessible exploration of filtered NMD AS events in iPSC-derived brain cell types, an interactive Shiny App was developed using the R packages *Shiny*, *Shinythemes*, and *ggplot2*. The database includes only NMD AS events meeting the criteria described above and contains information on gene expression levels, NMD AS event prediction information, PSI in the different cell types and splice junction read counts of reads supporting the AS event across all samples. To enhance context, gene expression data were extended with neuron expression profiles from comparable studies using the same iPSC line and *Ngn2*-based differentiation protocol at multiple developmental stages (DIV21, DIV28, and DIV49) (40–42).

### Conservation & GO-term analysis of common NMD events

To evaluate the conservation of common NMD AS events, the conservation of genomic regions flanking non-context dependent NMDin AS (mean PSI > 10%), non-context dependent NMDex AS (mean PSI < 90%) were compared to non-NMD AS events. Conservation was quantified using phastCons100way scores for each nucleotide position within 50 bp of the exon and 300 bp of the adjacent intron relative to the relevant splice junctions. Positions within introns extending into the next exon, or within exons extending into adjacent introns, were excluded. To minimize PSI-related bias between NMD AS and non-NMD AS events, NMDin AS and NMDex AS events were analyzed separately and compared to PSI-matched non-NMD AS event controls, generated using the *MatchIt* R package with the “nearest” method. Conservation differences were tested for each position and event-type (A3, A5, SE, RI, MX) using Student’s *t*-tests with p-values adjusted for multiple comparisons using the Benjamini Hochberg method. Due to the small number of events, NMDex AS retained intron events were excluded from analysis.

To assess whether genes harboring NMD AS events differ in evolutionary constraint compared to those without, we compared the pLI values of genes that contain non-context dependent NMD AS events to genes that do not. To minimize bias from NMD AS events being more frequently detected in highly covered genes, control genes were matched on long-read gene coverage using the *MatchIt* package (method = “optimal”). Differences in pLI distributions between groups were evaluated using the Wilcoxon rank-sum test.

To identify biological processes associated with NMD AS genes, we performed gene set enrichment analysis using *topGO*. To enrich for potential NMD AS events of biological relevance, only non-context dependent NMD AS events with significant CHX-DMSO PSI changes and >20% NMD-causing splicing were included. Control genes without NMD AS events were matched on long-read coverage as described above, and GO-term enrichment for biological processes was assessed using *topGO*.

### Identifying differentially spliced AS-events

To identify DS AS events a two-way ANOVA was performed on the PSI derived from CHX-treated samples of all four iPSC-derived cell types. *P*-values were corrected for multiple testing using the Benjamini-Hochberg method, and AS events were considered significantly differentially spliced if the adjusted *p* < 0.05 and the PSI difference was ≥ 0.2 between at least two samples. Context-dependent NMD AS events were excluded and only AS events with PSI available for all samples were included in the analysis. DS-NMDin and DS-NMDex AS events overlapping with DS-non-NMD AS events were removed using a graph-based approach, in which connected components of overlapping AS events (i.e., events fully or partially contained within another) were excluded. For significant AS events, a post hoc Tukey’s test was used to determine which specific cell-type comparisons contributed to the overall significance.

### Establishing inter-cell-type patterns for DS AS events

To identify AS events with shared inter-cell-type patterns, DS AS events were grouped by their unique combinations of significant pairwise comparisons, and inter-cell-type patterns were visualized using the ComplexUpset package in ‘distinct’ mode (43). For each inter-cell-type splicing pattern, gene set enrichment analysis was performed using topGO with Biological Process ontology, comparing all genes containing DS AS events within the respective inter-cell-type splicing pattern compared to non-DS AS events. The top 100 GO terms were selected irrespective of their significance, to account for patterns with few AS events. These GO terms were then clustered by shared gene members using the *simplifyEnrichment* R package. To assess the relative contribution of DS-NMD and DS-non-NMD AS events the ratio of DS-NMD or DS-non-NMD genes contributing to any GO term in that cluster was divided by the total number of DS-NMD or DS-non-NMD genes in the inter-cell-pattern, respectively.

### Identifying co-spliced AS events within inter-cell-type patterns

To characterize co-spliced AS events within each inter-cell-type pattern, DS AS events were clustered based on their relative average PSI across the CHX-treated conditions of the four different cell types. To obtain the relative average PSI, the average PSI was scaled from 0 (lowest average PSI) to 1 (highest) per AS event. Hierarchical clustering was performed on the relative average PSI using complete linkage. The optimal number of clusters (splicing profiles) was determined using the elbow method on within-cluster sum of squares (WSS): ranging between 2-4 clusters per inter-cell-type pattern.

### Exploring the effect of PTBP2 on DS AS events

To assess enrichment of PTBP2-regulated AS events within the clusters of co-spliced AS events, publicly available rMATS splicing data of skipped exon events following *PTBP2* knockdown in iPSC-derived neurons was used (7). Coordinates were converted from hg19 to hg38 using UCSC liftOver. skipped exon events from the top five inter-cell-type splicing patterns were then matched to skipped exon events in the *PTBP2*-KD data. AS events were considered significantly differentially spliced upon *PTBP2*-KD if adjusted *p* < 0.05 and absolute IncLevelDifference > 0.1. To reduce bias from higher detection of DS AS events in well-covered genes, genes containing DS AS events and genes without DS AS events were matched on total long-read coverage using the R package *MatchIt* (method = “optimal”). Enrichment of PTBP2-responsive DS AS events in each cluster within the inter-cell-type patterns was assessed with Fisher’s exact test and adjusted for multiple testing using the Benjamini Hochberg method. For clusters enriched in DS AS events that were also differentially spliced upon *PTBP2* KD, the distribution of differential gene expression between genes containing DS-NMD AS events and DS-non-NMD AS events upon *PTBP2* KD was compared using a chi-square test. Again, genes within the compared groups were matched on long-read gene coverage using *MatchIt*. Due to the low number of NMD AS events in cluster 1 of the glia-neuron inter-cell-type pattern, enrichment of differential expressed genes could not be assessed for this cluster.

### Selection of NMD AS events in NDD-associated genes

To identify NDDs that could potentially benefit from the TANGO approach, NMD AS events within NDD-associated genes were selected based on a previously published dataset of NDD genes annotated with ClinVar variants, disease inheritance and phenotypic feature information (9). Only NDD-genes related with a source score of >2 were included. To identify abundant NMD AS events, (context-dependent) NMDin AS events needed to have a PSI of higher than 0.1 and (context-dependent) NMDex AS events needed to have a PSI of lower than 0.9 in the CHX condition of at least one cell type. To remove false positive events and events with inflated PSI due to spurious annotations or overlapping AS events, all selected NMD AS events were manually evaluated on long-read transcript data and assessed for potential PSI bias resulting from AS events that share splice junctions. The molecular mechanism of all NDDs containing the remaining NMD AS events was manually annotated based on literature.

### Comparison with an existing NMD AS dataset

To compare our NMD AS dataset with a previously published dataset (8), all AS events in our unfiltered dataset were matched based on strand, event start, and event end coordinates. For each matched AS event found in the previously reported dataset, we assessed whether it was retained in our final filtered dataset. If the AS event was excluded the reason for exclusion was classified as: low relative event coverage (<10%), insufficient splice-context group support in the splice-context aware method (<2), AS event creating a long exon (>500 bp) or a combination of these factors. Abundant NMD AS events were defined as follows: (context-dependent) NMDin events with PSI > 0.1 and (context-dependent) NMDex events with PSI < 0.9 in the CHX condition of at least one cell type. To refine a systematic analysis of TANGO potential in NDDs, the presence of NMD AS events in NDD genes was reassessed in a previously published dataset with the previously published methodology using the *ComplexUpset* package with mode intersect (9, 43), but considering only NDDs with abundant NMD AS events (>10% PSI) in our NMD AS dataset.

## Acknowledgments

We would like to thank all members of the Tailored consortium for their invaluable support and feedback throughout this work. We thank Annelot van Esbroeck-Weevers and Raphael Kubler for the insightful input on the design and content of the Shiny application, as well as their constructive comments on the manuscript. We are grateful to everyone at the Radboud Genomics Technology Center and the Radboud Institute for Molecular Life Sciences sequencing facility for their collaborative input, support, and assistance in generating the long-read and short-read sequencing data. This study was supported by the Dutch Organisation for Health Research and Development (015.014.066 to LELMV), (10250022110002 to NNK and LELMV) and Simons Foundation (SFARI) Grant 890042 (to NKK). In addition, the aims of this study contribute to the Solve-RD project (to LELMV), and the ERDERA partnership program (to LELMV) which is supported by the European Union’s Horizon 2020 research and innovation program under grant agreements No. 779257 and No. 101156595, respectively. Views and opinions expressed are those of the author(s) only and do not necessarily reflect those of the European Union or any other granting authority, who cannot be held responsible for them. LELMV is a member of the European Reference Network on Rare Congenital Malformations and Rare Intellectual Disability ERN-ITHACA [EU Framework Partnership Agreement ID: 3HP-HPFPA ERN-01-2016/739516].

## References

1. Raj B, Blencowe BJ. Alternative Splicing in the Mammalian Nervous System: Recent Insights into Mechanisms and Functional Roles. Neuron. 2015;87(1):14–27.

2. Yan Q, Weyn-Vanhentenryck SM, Wu J, Sloan SA, Zhang Y, Chen K, et al. Systematic discovery of regulated and conserved alternative exons in the mammalian brain reveals NMD modulating chromatin regulators. Proc Natl Acad Sci U S A. 2015;112(11):3445–50.

3. Zheng S. Alternative splicing and nonsense-mediated mRNA decay enforce neural specific gene expression. Int J Dev Neurosci. 2016;55:102–8.

4. Zheng S, Gray EE, Chawla G, Porse BT, O’Dell TJ, Black DL. PSD-95 is post-transcriptionally repressed during early neural development by PTBP1 and PTBP2. Nat Neurosci. 2012;15(3):381–8, S1.

5. Makeyev EV, Zhang J, Carrasco MA, Maniatis T. The MicroRNA miR-124 promotes neuronal differentiation by triggering brain-specific alternative pre-mRNA splicing. Mol Cell. 2007;27(3):435–48.

6. Vuong JK, Lin CH, Zhang M, Chen L, Black DL, Zheng S. PTBP1 and PTBP2 Serve Both Specific and Redundant Functions in Neuronal Pre-mRNA Splicing. Cell Rep. 2016;17(10):2766–75.

7. Dawicki-McKenna JM, Felix AJ, Waxman EA, Cheng C, Amado DA, Ranum PT, et al. Mapping PTBP2 binding in human brain identifies SYNGAP1 as a target for therapeutic splice switching. Nat Commun. 2023;14(1):2628.

8. Lim KH, Han Z, Jeon HY, Kach J, Jing E, Weyn-Vanhentenryck S, et al. Antisense oligonucleotide modulation of non-productive alternative splicing upregulates gene expression. Nat Commun. 2020;11(1):3501.

9. Wijnant KN, Nadif Kasri N, Vissers L. Systematic analysis of genetic and phenotypic characteristics reveals antisense oligonucleotide therapy potential for one-third of neurodevelopmental disorders. Genome Med. 2025;17(1):59.

10. Han Z, Chen C, Christiansen A, Ji S, Lin Q, Anumonwo C, et al. Antisense oligonucleotides increase Scn1a expression and reduce seizures and SUDEP incidence in a mouse model of Dravet syndrome. Sci Transl Med. 2020;12(558).

11. Sullivan J, Wirrell E, Knupp KG, Chen D, Flamini R, Zafar M, et al. Adaptive functioning and neurodevelopment in patients with Dravet syndrome: 12-month interim analysis of the BUTTERFLY observational study. Epilepsy Behav. 2024;151:109604.

12. Yang R, Feng X, Arias-Cavieres A, Mitchell RM, Polo A, Hu K, et al. Upregulation of SYNGAP1 expression in mice and human neurons by redirecting alternative splicing. Neuron. 2023.

13. Spangsberg Petersen US, Dembic M, Martinez-Pizarro A, Richard E, Holm LL, Havelund JF, et al. Regulating PCCA gene expression by modulation of pseudoexon splicing patterns to rescue enzyme activity in propionic acidemia. Mol Ther Nucleic Acids. 2024;35(1):102101.

14. Mittal S, Tang I, Gleeson JG. Evaluating human mutation databases for “treatability” using patient-customized therapy. Med. 2022;3(11):740–59.

15. Pigini P, Xu H, Ji Y, Lindmeier H, Saltzman HR, Yun S, et al. Poison exon splicing in the human brain: a new paradigm for understanding and targeting neurological disorders. bioRxiv. 2025.

16. Frega M, van Gestel SH, Linda K, van der Raadt J, Keller J, Van Rhijn JR, et al. Rapid Neuronal Differentiation of Induced Pluripotent Stem Cells for Measuring Network Activity on Micro-electrode Arrays. J Vis Exp. 2017(119).

17. Zhang Y, Pak C, Han Y, Ahlenius H, Zhang Z, Chanda S, et al. Rapid single-step induction of functional neurons from human pluripotent stem cells. Neuron. 2013;78(5):785–98.

18. Schuurmans IME, Mordelt A, Linda K, Puvogel S, Duineveld D, Hommersom MP, et al. Navigating Human Astrocyte Differentiation: Direct and Rapid one-step Differentiation of Induced Pluripotent Stem Cells to Functional Astrocytes Supporting Neuronal Network development. bioRxiv. 2024:2024.03.27.586938.

19. Mordelt A, Schuurmans IME, Scheefhals N, Hommersom MP, Slottje K, Mast K, et al. Long-term co-maturation of stem cell-derived microglia and neuronal networks: an optimized platform to assess human microglial contribution to neuronal function. bioRxiv. 2025:2025.10.24.684345.

20. Zhou H, Deng XW. Intron Retention, an Orchestrated Program of Gene Expression Regulation. Bioessays. 2025;47(4):e202400248.

21. Green ID, Pinello N, Song R, Lee Q, Halstead JM, Kwok CT, et al. Macrophage development and activation involve coordinated intron retention in key inflammatory regulators. Nucleic Acids Res. 2020;48(12):6513–29.

22. Ni JZ, Grate L, Donohue JP, Preston C, Nobida N, O’Brien G, et al. Ultraconserved elements are associated with homeostatic control of splicing regulators by alternative splicing and nonsense-mediated decay. Genes Dev. 2007;21(6):708–18.

23. Zhu B, Fisher E, Li L, Zhong P, Yan Z, Feng J. PTBP2 attenuation facilitates fibroblast to neuron conversion by promoting alternative splicing of neuronal genes. Stem Cell Reports. 2023;18(11):2268–82.

24. Akter M, Cui H, Hosain MA, Liu J, Duan Y, Ding B. RANBP17 Overexpression Restores Nucleocytoplasmic Transport and Ameliorates Neurodevelopment in Induced DYT1 Dystonia Motor Neurons. J Neurosci. 2024;44(15).

25. Pecoraro S, Verkerke M, Sluijs JA, van Het Hof B, van der Pol SMA, van Strien ME, et al. Adult human subventricular zone microglia promote a pro-neurogenic niche for neuronal progenitors in Parkinson’s disease. Brain Behav Immun. 2025;129:318–34.

26. He Y, Wang Y, Yu H, Tian Y, Chen X, Chen C, et al. Protective effect of Nr4a2 (Nurr1) against LPS-induced depressive-like behaviors via regulating activity of microglia and CamkII neurons in anterior cingulate cortex. Pharmacol Res. 2023;191:106717.

27. Hu D, Huang C, Tang L, Lei J, Wang J, Hu W, et al. NR4A2 attenuates early brain injury after intracerebral hemorrhage by promoting M2 microglial polarization via TLR4/TRAF6/NF-kappaB pathway inhibition. Cell Mol Life Sci. 2025;82(1):262.

28. Zhang Y, Chen K, Sloan SA, Bennett ML, Scholze AR, O’Keeffe S, et al. An RNA-sequencing transcriptome and splicing database of glia, neurons, and vascular cells of the cerebral cortex. J Neurosci. 2014;34(36):11929–47.

29. Su CH, D D, Tarn WY. Alternative Splicing in Neurogenesis and Brain Development. Front Mol Biosci. 2018;5:12.

30. Cai W, Dai X, Chen J, Zhao J, Xu M, Zhang L, et al. STAT6/Arg1 promotes microglia/macrophage efferocytosis and inflammation resolution in stroke mice. JCI Insight. 2019;4(20).

31. Herculano-Houzel S. The glia/neuron ratio: how it varies uniformly across brain structures and species and what that means for brain physiology and evolution. Glia. 2014;62(9):1377–91.

32. Sardar D, Lozzi B, Woo J, Huang TW, Cvetkovic C, Creighton CJ, et al. Mapping Astrocyte Transcriptional Signatures in Response to Neuroactive Compounds. Int J Mol Sci. 2021;22(8).

33. Gosselin D, Skola D, Coufal NG, Holtman IR, Schlachetzki JCM, Sajti E, et al. An environment-dependent transcriptional network specifies human microglia identity. Science. 2017;356(6344).

34. Dobin A, Davis CA, Schlesinger F, Drenkow J, Zaleski C, Jha S, et al. STAR: ultrafast universal RNA-seq aligner. Bioinformatics. 2013;29(1):15–21.

35. Kuo RI, Cheng Y, Zhang R, Brown JWS, Smith J, Archibald AL, et al. Illuminating the dark side of the human transcriptome with long read transcript sequencing. BMC Genomics. 2020;21(1):751.

36. Tang S, Lomsadze A, Borodovsky M. Identification of protein coding regions in RNA transcripts. Nucleic Acids Res. 2015;43(12):e78.

37. Pardo-Palacios FJ, Arzalluz-Luque A, Kondratova L, Salguero P, Mestre-Tomas J, Amorin R, et al. SQANTI3: curation of long-read transcriptomes for accurate identification of known and novel isoforms. Nat Methods. 2024;21(5):793–7.

38. Trincado JL, Entizne JC, Hysenaj G, Singh B, Skalic M, Elliott DJ, et al. SUPPA2: fast, accurate, and uncertainty-aware differential splicing analysis across multiple conditions. Genome Biol. 2018;19(1):40.

39. Wang Y, Xie Z, Kutschera E, Adams JI, Kadash-Edmondson KE, Xing Y. rMATS-turbo: an efficient and flexible computational tool for alternative splicing analysis of large-scale RNA-seq data. Nat Protoc. 2024;19(4):1083–104.

40. Yuan X, Puvogel S, van Rhijn JR, Ciptasari U, Esteve-Codina A, Meijer M, et al. A human in vitro neuronal model for studying homeostatic plasticity at the network level. Stem Cell Reports. 2023;18(11):2222–39.

41. Hommersom MP, Doorn N, Puvogel S, Lewerissa EI, Mordelt A, Ciptasari U, et al. CACNA1A haploinsufficiency leads to reduced synaptic function and increased intrinsic excitability. Brain. 2025;148(4):1286–301.

42. Leonardi O, Lewerissa E, Aprile D, Barakat TS, Culotta L, Deng R, et al. CHD2 Dosage Ties Autolysosomal Pathway to Cortical Maturation in Disease and Evolution. bioRxiv. 2025:2025.01.21.634145.

43. Lex A, Gehlenborg N, Strobelt H, Vuillemot R, Pfister H. UpSet: Visualization of Intersecting Sets. IEEE Trans Vis Comput Graph. 2014;20(12):1983–92.

