## Supplementary Figures for "Mapping NMD-coupled alternative splicing in iPSC-derived brain cells: a resource for therapeutic discovery"

A.

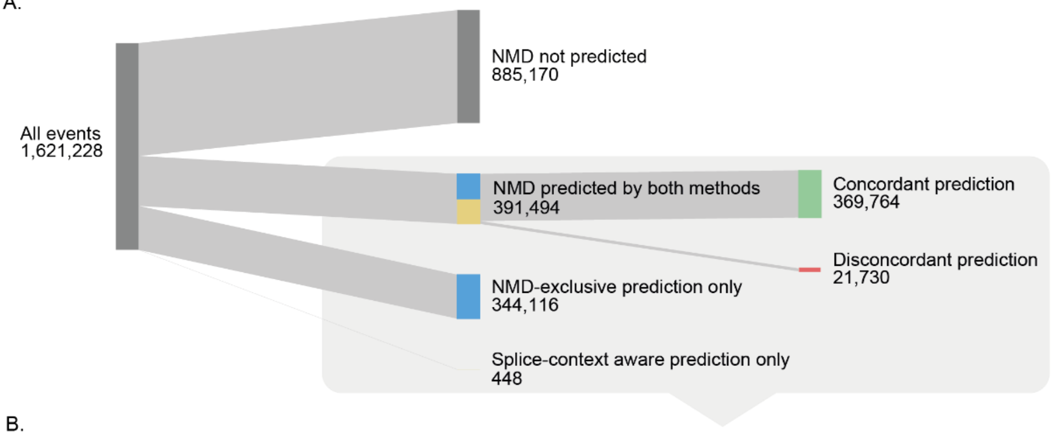


B.

| NMD exclusive | Splice-context aware | | | | | | | |
|---|---|---|---|---|---|---|---|---|
| | *nonNMD* | *NMDin* | *NMDex* | *NMDin_NMDex* | *NMDin_nonNMD* | *NMDex_nonNMD* | *undetermined* | *inconclusive* |
| *nonNMD* | 218359 | 11779 | 4470 | 518 | 2716 | 2247 | 139484 | 3067 |
| *NMDin* | 0 | 84633 | 0 | 0 | 0 | 0 | 49485 | 13597 |
| *NMDex* | 0 | 0 | 66772 | 0 | 0 | 0 | 43737 | 11309 |
| *NMDin_NMDex* | 0 | 0 | 0 | 0 | 0 | 0 | 35509 | 47928 |
| *not_in_CDS* | 448 | 0 | 0 | 0 | 0 | 0 | 0 | 0 |

***Supplementary Figure 1. Overview of predicted AS events by two NMD AS prediction methods. A.*** *Sankey's diagram of NMD AS prediction of all AS events demonstrating the number of AS events predicted by the NMD-exclusive method and splice-context aware method.* ***B.*** *Table of NMD AS prediction of all unfiltered AS events predicted by at least one of two NMD-prediction methods (highlighted by a grey square in panel A). Background colors correspond to the nodes in the Sankey's diagram: concordant events (receiving the same prediction using both methods, green), only predicted by the NMD-exclusive method (blue), only predicted by the splice-context aware method (yellow) or disconcordant (receiving different predictions using both methods, red). NMD = nonsense-mediated decay, NMDin = AS events predicted to cause NMD upon inclusion, NMDex = AS events predicted to cause NMD upon exclusion, non-NMD = AS events not predicted to cause NMD.*

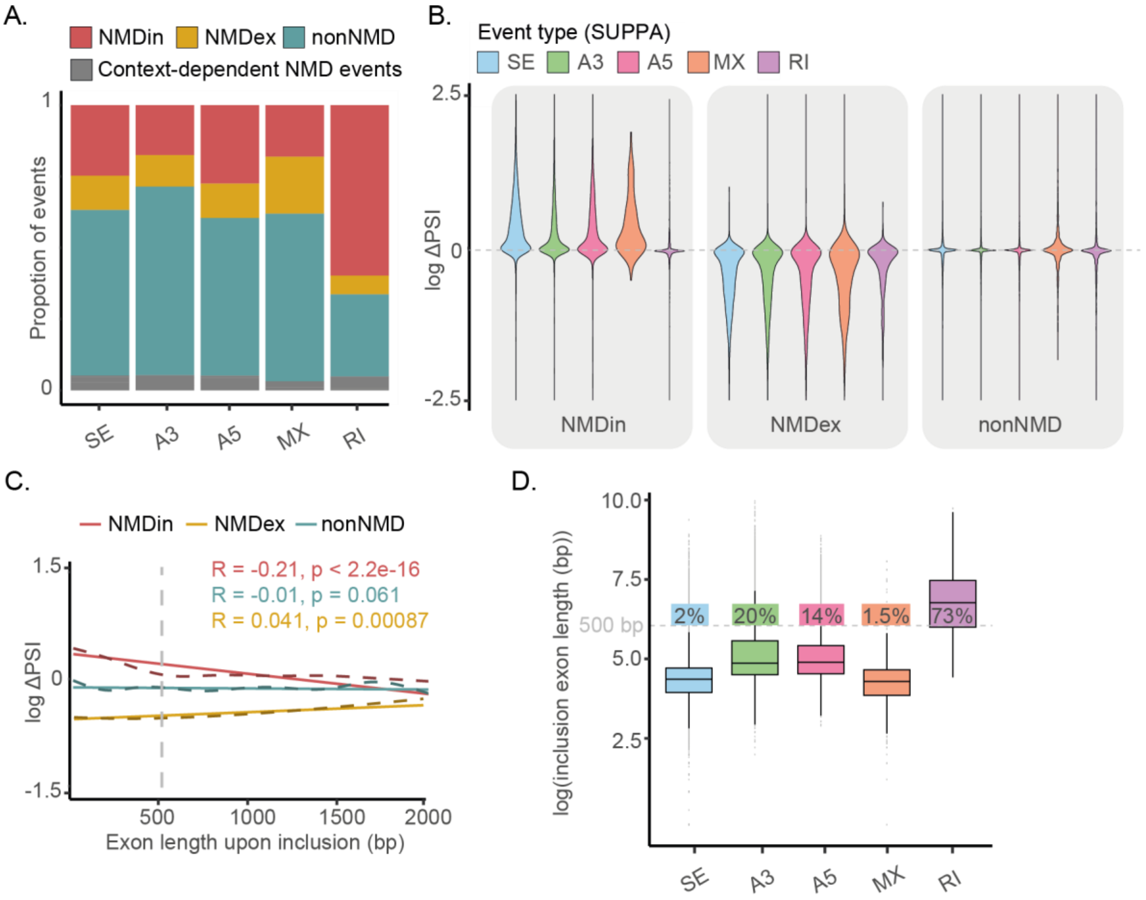


***Supplementary Figure 2. Intron-retentions and AS events creating long-exons are NMD resistant.*** ***A.*** *The proportion of NMD types across different SUPPA AS event categories.* ***B.*** *Violin plots of log ΔPSI (PSI in CHX condition - PSI in DMSO control condition) per NMD type separated by SUPPA AS event-types with NMDin and NMDex AS events showing typical skewing towards a positive and negative log ΔPSI, respectively, for all SUPPA AS event-types except NMDin retained intron events.* ***C.*** *Correlation between log ΔPSI (PSI in CHX condition - PSI in DMSO control condition) and exon length upon AS event inclusion; calculated using Pearson correlation.* Sample sizes: NMDin (n = 15,391), NMDex (n = 6,750), nonNMD (n = 33,518). ***D.*** *Distribution of log exon lengths after AS event inclusion for each SUPPA AS event category; the proportion of AS events >500 bp, per SUPPA AS event category is indicated in the plot. * p < 0.05, ** p < 0.01, *** p < 0.001 and **** p < 0.0001. PSI = percentage spliced in, A3 = alternative 3’ splice site, A5 = alternative 5’ splice site, MX = mutually exclusive, RI = retained intron, SE = skipped exon, NMD = nonsense-mediated decay, NMDin = AS events predicted to cause NMD upon inclusion, NMDex = AS events predicted to cause NMD upon exclusion, non-NMD = AS events not predicted to cause NMD.*

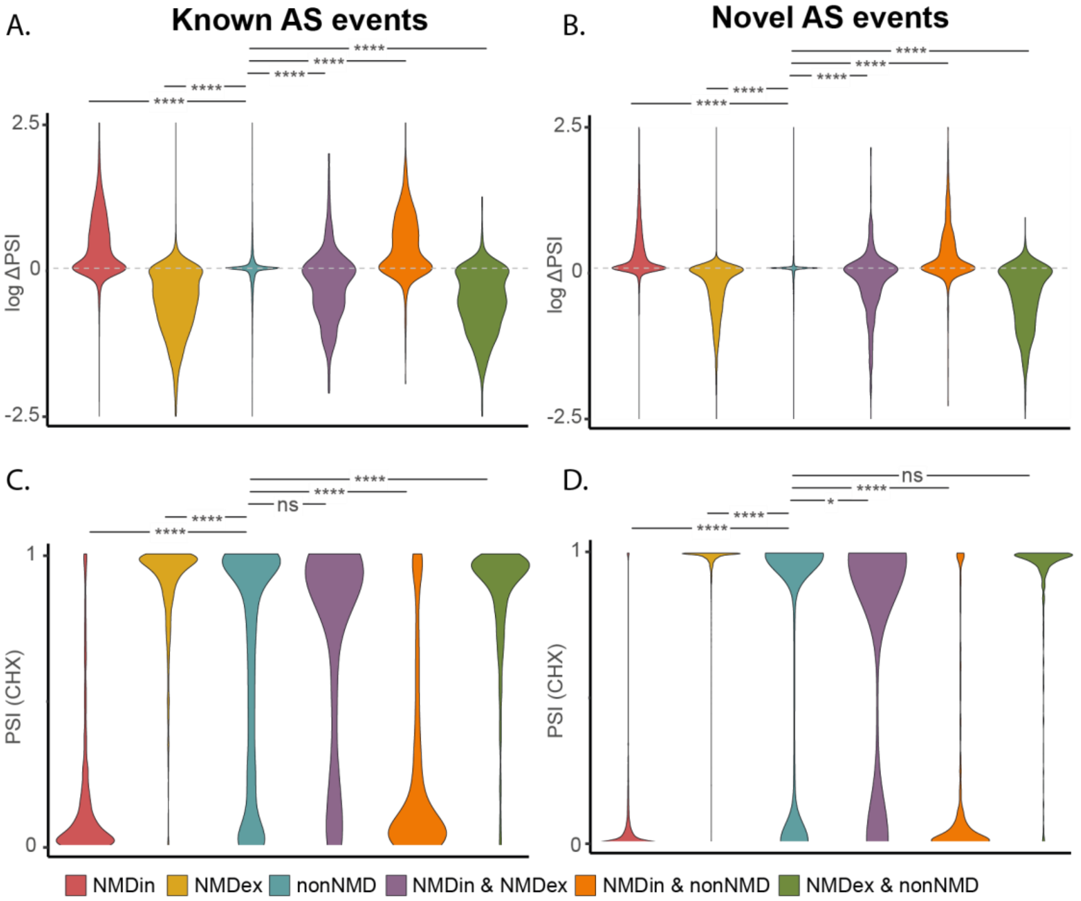


***Supplementary Figure 3. PSI and ΔPSI of novel and known AS events A/B.*** *Violin plots of log ΔPSI (PSI in CHX condition - PSI in DMSO control condition) per NMD AS type of known AS events (**A**) and novel AS events (**B**).* ***C/D.*** *Violin plots of mean CHX-treated PSI value distribution per NMD AS type of known AS events (**C**) and novel AS events (**D**). * p < 0.05, ** p < 0.01, *** p < 0.001 and **** p < 0.0001. NMD = nonsense-mediated decay, NMDin = AS events predicted to cause NMD upon inclusion, NMDex = AS events predicted to cause NMD upon exclusion, non-NMD = AS events not predicted to cause NMD, PSI = percentage spliced in.*

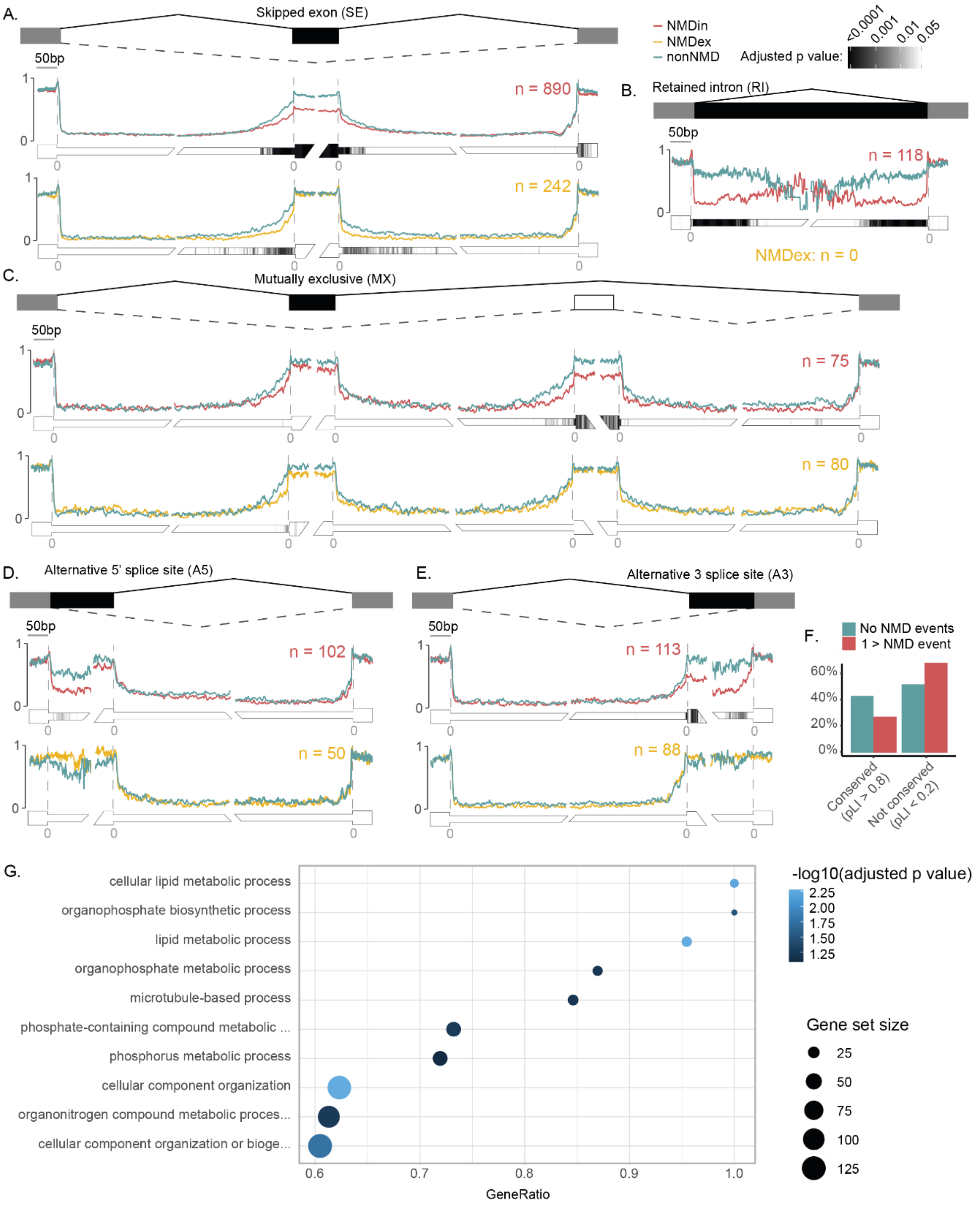


***Supplementary Figure 4. Conservation and functional enrichment of NMD AS events. A/B/C/D/E.*** *Average conservation profiles around splice junctions of alternative splicing events. Each plot shows the mean PhastCons100 score averaged over all AS events of the indicated type on the y-axis (representing the average evolutionary conservation across 100 vertebrate genomes, 0 = no conservation, 1 = fully conserved) at each position relative to the splice site on the x axis. Conservation is shown for intronic and exonic regions flanking splice junctions of NMDin AS events (red), NMDex AS events (yellow) and non-NMD AS events (blue), for different SUPPA AS event types; (**A**) skipped exons (SE), (**B**) retained introns (RI), (**C**) mutually exclusive exons (MX), (**D**) alternative 5' splice sites (A5), and (**E**) alternative 3' splice sites (A3).* ***F.*** *Percentage of genes within each NMD group (no NMD AS events, blue; more than one NMD AS event, red) that are conserved (pLI > 0.8) or*

*non-conserved (pLI < 0.2).* ***G.*** *Top 10 GO-terms of genes containing common NMD AS events (>20% NMD). NMD = nonsense-mediated decay. NMD = nonsense-mediated decay, NMDin = AS events predicted to cause NMD upon inclusion, NMDex = AS events predicted to cause NMD upon exclusion, non-NMD = AS events not predicted to cause NMD.*

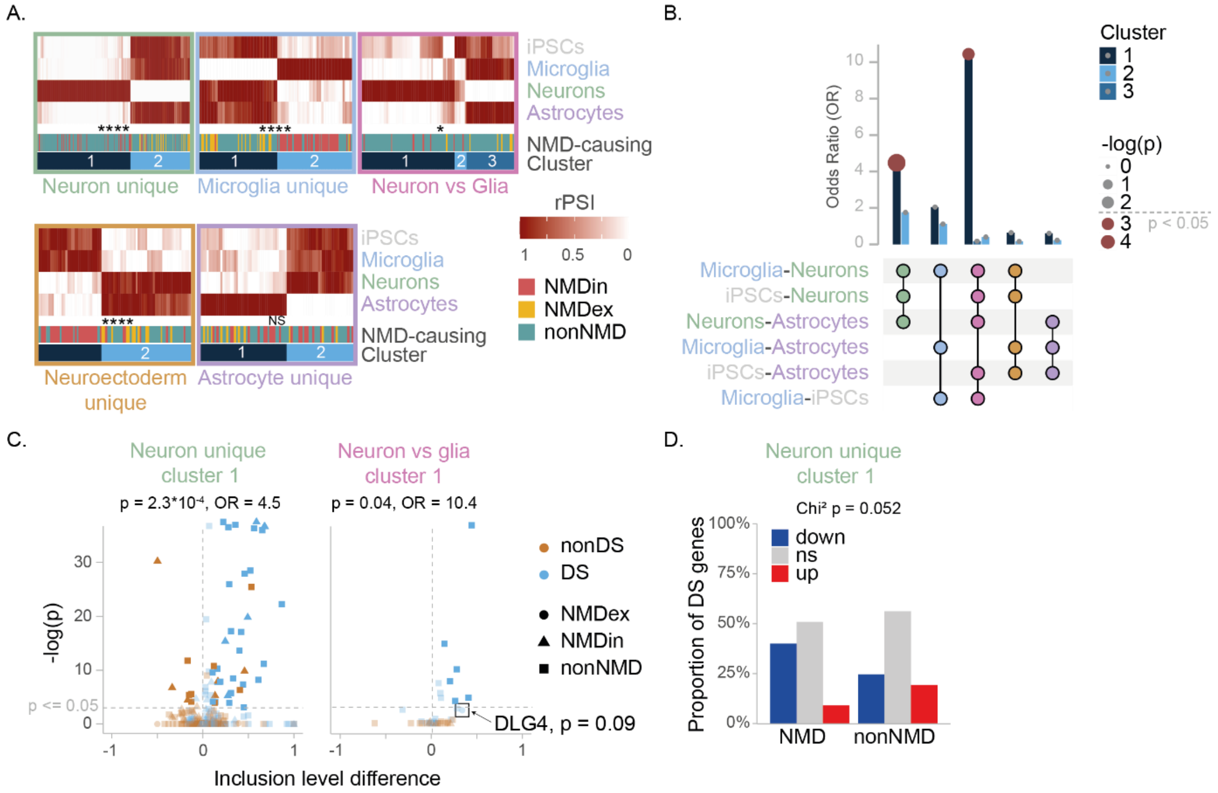


***Supplementary Figure 5. Knock down of PTBP2 affects clusters of differentially spliced AS events. A.** Heatmap of relative PSI (rPSI, scaled 0-1 per event, with the lowest PSI set to 0 and the highest to 1), annotated for NMD AS event type (non-NMD, NMDin and NMDex) and cluster membership within the top 5 inter-cell-type patterns. Asterixes indicate a significant shift in the distribution of NMDin versus NMDex events across AS event clusters within each inter-cell-type pattern, calculated using a chi-square test. **B.** Upset plot of significant pairwise comparisons (Tukey) of DS AS-events, representing the top 5 inter-cell-type patterns. Combinations of significant comparisons are indicated by connected dots with a barplot representing the odds ratio (OR) and significance of DS events upon PTBP2 knockdown for each cluster (corresponding to panel A) within the inter-cell-type patterns. **C.** Volcano plot of differential splicing in a PTBP2 knockdown dataset for DS AS events (blue) and PSI-matched non-DS event (orange) identified from our dataset for two selected clusters (corresponding to panel A). **D.** Distribution of significantly down- and upregulated genes upon PTBP2 knockdown for DS NMD AS event containing genes versus DS non-NMD AS event containing genes within cluster 1 of the neuron-unique inter-cell-type splice pattern. * $p < 0.05$, ** $p < 0.01$, *** $p < 0.001$, and **** $p < 0.0001$. NMD = nonsense mediated decay, PSI = percentage spliced in, NMDin = AS events predicted to cause NMD upon inclusion, NMDex = AS events predicted to cause NMD upon exclusion, non-NMD = AS events not predicted to cause NMD.*

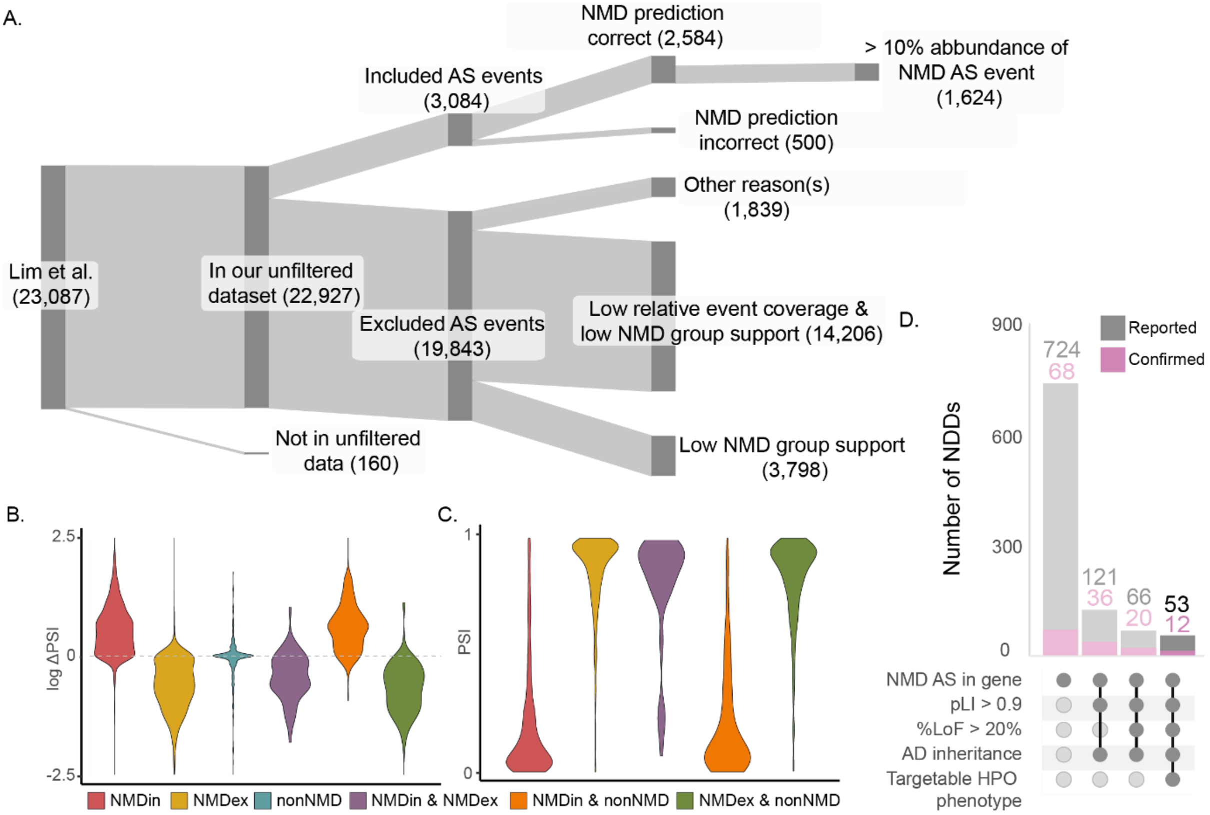


***Supplementary Figure 6. Comparison of our NMD AS database with reported NMD AS events. A.*** *Sankey diagram illustrating the fate of NMD AS events reported by Lim et al.(8) in our NMD AS database, showing if each AS event is present in the unfiltered dataset, if it is included or excluded, and if it is predicted to induce NMD in our dataset.* ***B.*** *Violin plot of ΔPSI (PSI in CHX condition - PSI in DMSO control condition) for NMD AS events reported by Lim et al. that are included in our NMD AS database, stratified by NMD AS event type (NMDin, NMDex, non-NMD, NMDin&NMDex, NMDin&non-NMD, NMDex&non-NMD), as predicted by the splice-context aware method. C. Violin plot of PSI for the same included NMD AS events, stratified by NMD AS event type, as predicted by the splice-context aware method.* ***D.*** *UpSet plot of favorable combinations of characteristics for TANGO for NDDs, adapted from Wijnant et al (9). The bar representing the most favorable combination of characteristics is highlighted. LoF = loss-of-function, pLI = probability of being loss-of-function intolerant, AD = autosomal dominant, HPO = Human phenotype ontology, NMD = nonsense-mediated decay, NMDin = AS events predicted to cause NMD upon inclusion, NMDex = AS events predicted to cause NMD upon exclusion, non-NMD = AS events not predicted to cause NMD.*

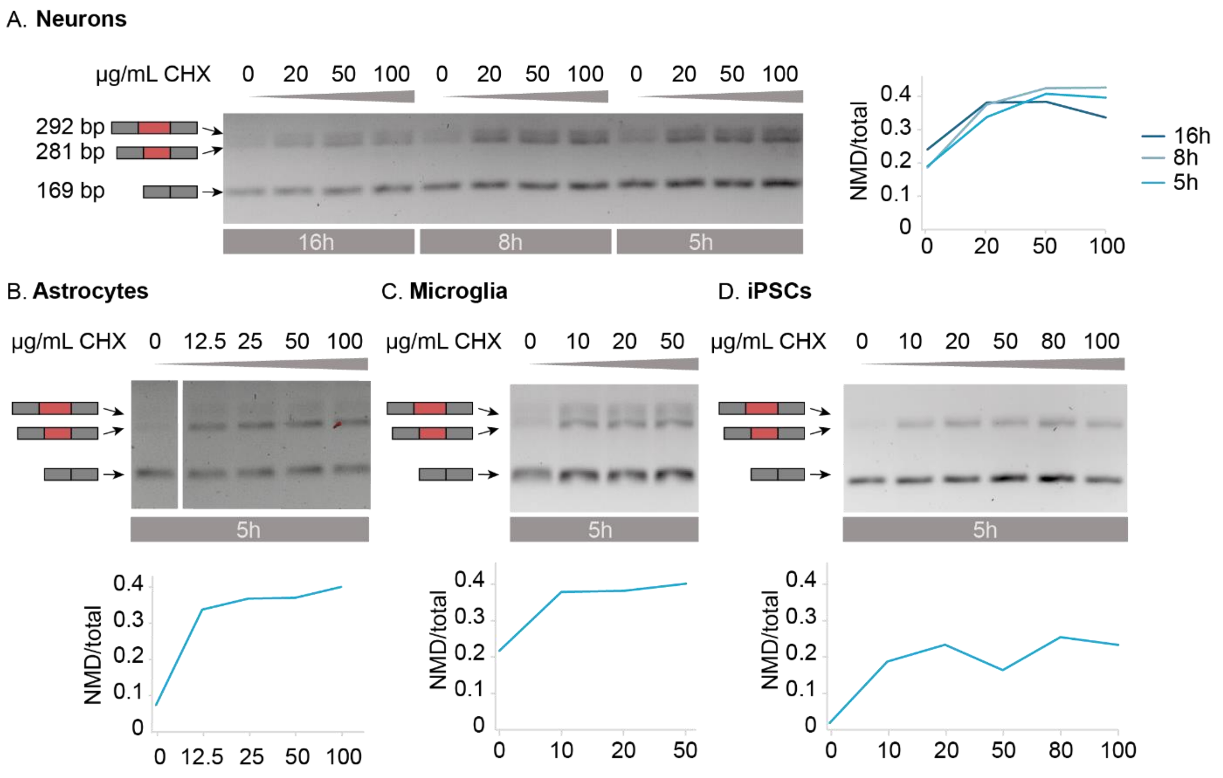


***Supplementary Figure 7. Optimization of CHX concentration and duration for NMD inhibition. A.*** *RT-PCR agarose gel of an NMD-in AS event in CHD2 across increasing CHX concentrations and durations in neurons. The 192 and 181 bp bands represent the NMD-sensitive inclusion isoform, and the 169 bp band represents the NMD-insensitive exclusion isoform. NMDin inclusion was quantified as the ratio of inclusion band intensity (192 + 181 bp) to the total band intensity (192 + 181 + 169 bp) and visualized as a line graph.* ***B/C/D.*** *RT-PCR agarose gel and NMDin inclusion quantification of an NMD-in AS event in CHD2 across increasing CHX concentrations in astrocytes, microglia and iPSCs. CHX = cycloheximide.*

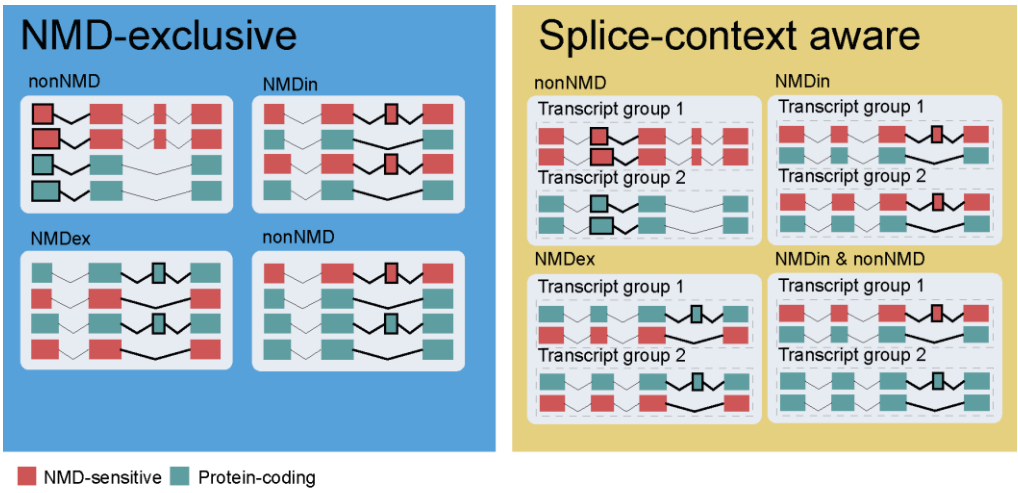


***Supplementary Figure 8. NMD-prediction methods.*** *Graphical overview of the two NMD AS event identification methods used: NMD-exclusive and splice-context aware.*
